# Identification of Obelisk-like covalently closed circular RNA replicon in hot springs by double-stranded RNA sequencing and expansion of the diversity of the Obelisk superfamily

**DOI:** 10.1101/2025.09.03.673927

**Authors:** Syun-ichi Urayama, Akihito Fukudome, Pascal Mutz, Yosuke Matsushita, Yoshihiro Takaki, Yosuke Nishimura, Sofia Medvedeva, Mart Krupovic, Eugene V. Koonin, Takuro Nunoura

## Abstract

Extensive metatranscriptome mining has recently vastly expanded the range of covalently closed circular (ccc) RNA replicons. A notable group of such replicons are Obelisks, cccRNAs of about 1 kilobase (kb) encoding a protein with a unique fold, Oblin-1, and detected in a broad variety of metatranscriptomes, in particular, those from the human gastrointestinal tract. We adopted Fragmented and primer-Ligated DsRNA Sequencing (FLDS) method to selectively sequence double-stranded (ds) RNAs, the replicative intermediates of RNA replicons, and to identify cccRNAs among the resulting sequences. From these data, we selected cccRNAs with predicted extensive intramolecular base-pairing, a hallmark of viroid-like elements. ch We employed FLDS to explore metatranscriptomes from acidic hot springs in Japan and discovered a distinct family of Obelisks probably associated with thermoacidophilic bacteria (Hot spring Obelisks, HsObs). The proteins encoded by HsObs, HsOblins, show no significant sequence similarity to previously identified Oblin-1 proteins, but are predicted to adopt a closely similar structure. A comprehensive search of metagenomes for Oblin-1 and HsOblin homologs substantially expanded this family of Obelisk-encoded proteins revealing several distinct subfamilies that share the same core fold. A cccRNA encoding an HsOblin homolog was also detected in a Yellowstone hot spring metatranscriptome. Apart from Oblin-1, some subfamilies of Obelisks were predicted to encode additional small proteins with simple alpha-helical folds.

## Introduction

Recently, a broad variety of covalently closed circular (ccc) RNAs have been identified in diverse cells, leading to increased interest in their origins and functions (Lasda and Parker, 2014; Liu and Chen, 2022; Navarro and Turina, 2024). Some of these cccRNAs originate from cellular genomes whereas other represent RNA replicons including viroids, viroid-like satellite RNAs and several groups of viruses (Forgia et al., 2023; Koonin et al., 2024; Koonin and Lee, 2025; Lee and Koonin, 2022). Viroids are the smallest and simplest known replicators, and the smallest known pathogens as well. Until recently, viroids have been identified only in a few plant species, in some of which they cause disease. The genomes of viroids are cccRNAs of only 200–400 nucleotides (nt) in size that form extensive rod-shaped or branched secondary structure and encode no proteins (Di Serio et al., 2014). Viroids fully depend on the host cell enzymatic machinery for their replication and are replicated by DNA-dependent RNA polymerases (RNAP) via a version of the rolling circle replication (RCR) mechanism (Flores et al., 2009). In one of the recognized families of viroids, *Avsunviroidae*, concatemeric RCR intermediates are processed by hammerhead ribozymes (HHR) present in both polarities of the viroid RNA (Di Serio et al., 2018). Viroids of the other family, *Pospiviroidae*, lack ribozymes, and the RCR intermediates are processed by host RNases (Di Serio et al., 2021).

Apart from the bona fide viroids, several groups of viroid-like cccRNA replicons have been identified. Satellite cccRNAs, sometimes referred to as virusoids, are similar to viroids but depend on plant RNA viruses, being replicated by the viral RNA-dependent RNA polymerase (RdRP) and packaged in the helper virus particles. Retroviroids are viroid-like cccRNA (CarSV) that also contain HHRs, but differ from viroids in that they are transcribed into a DNA form by reverse transcriptases of helper plant pararetroviruses (family *Caulimoviridae*) and can integrate into the plant genomic DNA. Retrozymes are small, nonautonomous retrotransposons identified in plant and animal genomes that depend on Ty3-like retrotransposons (family *Metaviridae*) for reverse transcription into a DNA form and integration into the host genome. The retrozyme transcripts are circularized by HHRs present in their long terminal repeats.

In addition to these non-coding viroid-like cccRNA replicons, an important human pathogen, hepatitis delta virus (HDV; family *Kolmioviridae*), a satellite of hepatitis B virus (HBV), has a cccRNA genome of about 1.7 kb that encodes a virus nucleocapsid protein (known as Hepatitis Delta Antigen, HDAg) and contains a unique ribozyme in both RNA polarities (Kuhn et al., 2024a). Like viroids, HDV genome is replicated by the host RNAP, whereas HBV provides the virion envelope protein. Until recently, HDV remained the only known viroid-like virus, but in the last few years, related viruses have been discovered in other vertebrates and arthropods. Because HDV and its relatives are unrelated to any other known viruses, the International Committee on Taxonomy of Viruses (ICTV) classified this unusual group of viruses into a separate viral realm (the highest rank in the taxonomy of viruses), *Ribozyviria*.

Recent efforts on metatranscriptome mining have transformed our understanding of the diversity and provenance of cccRNA replicons (e.g. (Koonin and Lee, 2025)). Thousands of distinct non-coding, ribozyme-containing cccRNAs in the viroid size range have been identified in metatranscriptomes from diverse environments many of which are virtually devoid of plant or animal material implying that these cccRNA replicons are hosted by unicellular eukaryotes and/or prokaryotes. Furthermore, several of these cccRNAs were found to be targeted by CRISPR systems in genomes of bacteria, strongly suggestive of bacterial hosts. In addition to these viroid-like cccRNAs, metatranscriptome mining led to the identification of several groups of viruses with cccRNA genomes (Lee et al., 2023). Many of these are diverse members of *Ribozyviria* encoding distant homologs of HDAg, whereas others are unrelated to ribozyviruses. A particularly notable group are ambiviruses that possess the largest known cccRNA genomes at nearly 5 kb and encode two proteins one of which is a distinct RdRP whereas the other one has no detectable homologs among known proteins. Some of the ambiviruses were previously identified in basidiomycete fungi but the cccRNA structure of their genomes has not been initially recognized (Chong and Lauber, 2023; Forgia et al., 2023; Lee et al., 2023). Because ambiviruses encode an RdRP, the hallmark of the kingdom *Orthornavirae* within the viral realm *Riboviria*, that shows no specific affinity to any other RdRPs and given their unique genome organization, ambiviruses are currently classified as phylum *Ambiviricota* within *Orthornavirae* (Kuhn et al., 2024b).

The most recent major discovery in the expanding domain of cccRNA replicons are Obelisks, cccRNAs of about 1 kb in size encoding a protein without detectable homologs denoted Oblin-1 (Zheludev et al., 2024). Obelisks were originally identified in the human gut metatranscriptomes but subsequently found in a broad variety of environments and appear to be particularly abundant in marine ecosystems (Lopez-Simon et al., 2025). For one of the Obelisks, the bacterium *Streptococcus sanguinis* has been definitively identified as the host (Zheludev et al., 2024).

In this work, we explored dsRNA metatranscriptomes constructed by Fragmented and primer-Ligated DsRNA Sequencing (FLDS) (Hirai et al., 2021; Urayama et al., 2016), a method for selectively sequencing of dsRNA, a typical replicative intermediates of RNA replicons (Morris and Dodds, 1979; Roossinck et al., 2010; Semancik, 1986). We searched the FLDS data for cccRNAs with predicted extensive intramolecular base-pairing, a hallmark of viroid-like elements. In dsRNA metatranscriptomes from geothermal acidic hot springs in Japan, some of which harbor RNA virus genomes (Urayama et al., 2024), we identified a distinct family of Obelisks. A comprehensive search for Oblin-1 homologs substantially expanded the diversity of the Obelisk superfamily of cccRNA replicons, notably revealing their presence in high-temperature environments beyond previously studied gastrointestinal and aquatic ecosystems.

## Main text

### Identification of circular RNAs from a dsRNA-seq transcriptome

The dsRNA-seq data include reads originating from cccRNA replicons due to their RCR replication via dsRNA intermediates (Branch and Robertson, 1984; Fall et al., 2020; Flores et al., 2009) and possibly matured cccRNAs with extensive intramolecular base-pairing (Semancik, 1986). We established a data processing workflow to identify candidate cccRNA replicons from FLDS data in a sequence similarity-independent manner (Fig. 1). As a test case, plant leaves harboring a known viroid, Chrysanthemum stunt viroid (CSVd, species *Pospiviroid impedichrysanthemi*), were subjected to FLDS. Of the 6,311 contigs generated by *de novo* assembly of clean FLDS reads using SPAdes (metaplasmid mode), 7 contigs were predicted by the ccfind program to be cccRNAs, based on terminal redundancy. Based on the BLASTX and BLASTN search results, three of these sequences were discarded because they showed significant similarity to known plant and human sequences. Two additional sequences were removed because of their low coverage (<10 average coverage). The two remaining sequences with more than 10× average coverage were predicted to form an extensive secondary structure (>60% paired bases). Using read mapping data, the normalized coefficient of variation (CV) of coverage and normalized entropy of read start point were calculated to evaluate evenness of mapping (Table S1). The BLASTN search showed that these two sequences were variants of CSVd, with more than 95% nucleotide identity to the CSVd reference sequence (accession: X16408) (Fig. 1). These results indicate that the combination of FLDS and the developed workflow with the appropriate filtering, enables the structure-dependent identification of circular RNA replicons.

**Fig. 1:**
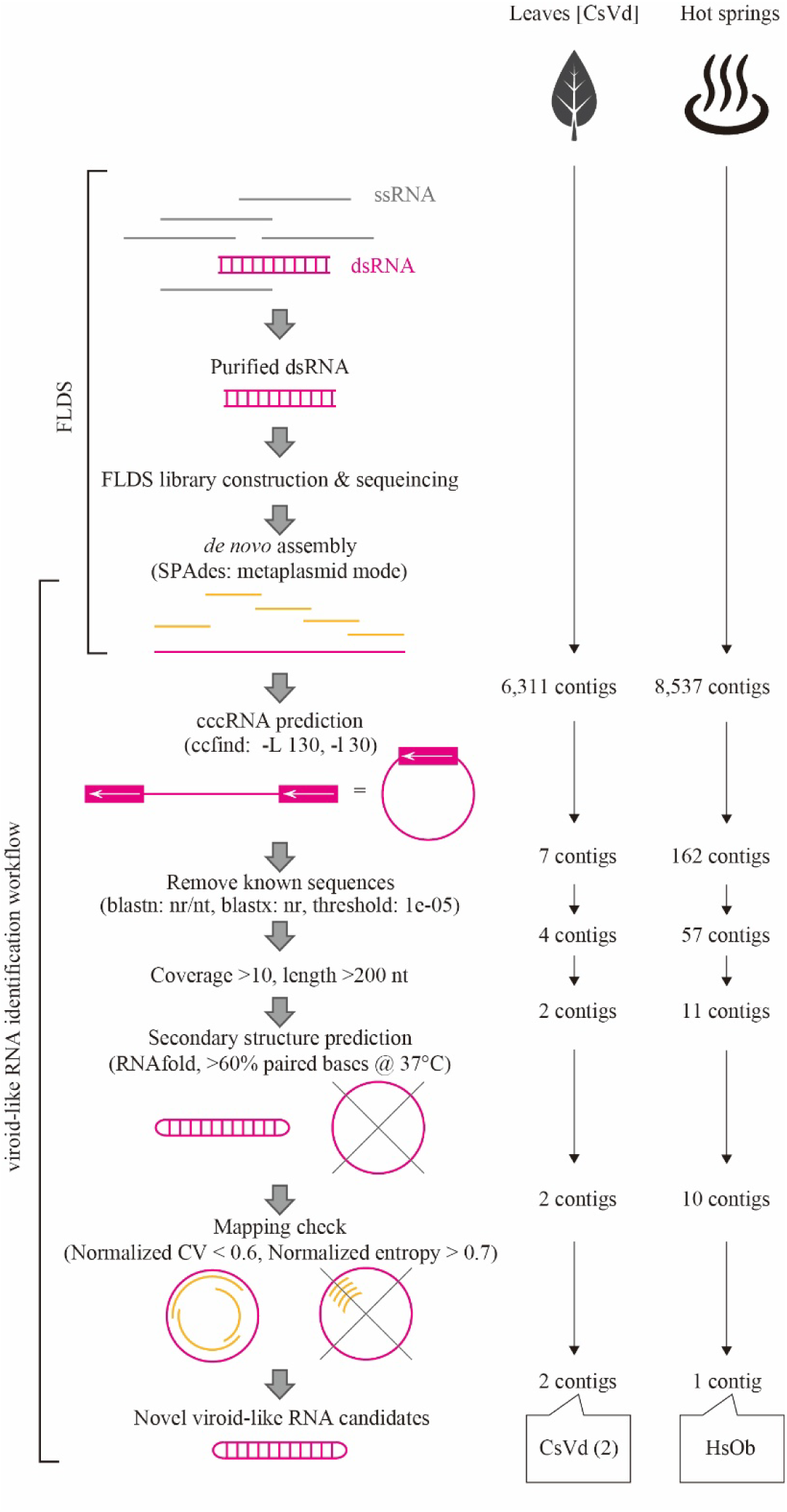
Schematic workflow of cccRNA replicon surveillance. DsRNAs obtained from samples were sequenced using the FLDS method (Urayama et al., 2024). The contigs assembled through *de novo* assembly were used for circular RNA prediction. After removing sequences with low coverage, those that were too short, and duplicated sequences, the predicted secondary structure and mapping patterns were also checked. Details of this workflow are described in the Materials and Methods section.

### Circular RNAs from acidic geothermal spring dsRNA metatranscriptomes

Using the workflow described above, a cccRNA candidate was identified from high-temperature acidic geothermal hot spring FLDS metatranscriptomes (Fig. 1, Table 1). Analysis with the ccfind program of 8,537 contigs obtained from the *de novo* assembly of metatranscriptomes using SPAdes for each sample yielded 162 candidate cccRNAs. Among these sequences, 105 were discarded because they showed significant similarity (*E*-value ≤ 1 × 10^−05^) to known non-cccRNA sequences in NCBI nr/nt or nr (protein) databases using BLASTN and BLASTX, respectively. After discarding short sequences (< 200 nt) and those with low average sequencing coverage (< 10), 11 candidate cccRNAs remained (Table S1). Of these, 10 were predicted to form a viroid-like secondary structure using RNAfold (>60% paired bases). Nine of these sequences were excluded from further consideration due to uneven read mapping, as determined by their normalized CV and entropy values (Table S1). Finally, a candidate sequence, Hot spring Obelisk (HsOb) (see below), was distinct.

**Table 1:** Characteristics of hot spring water samples.

| Code | Geographical<br>coordination | Area | Temp<br>(°C) | pH | DO<br>(mg/L) | H<sub>2</sub>S<br>(mM) | Sampling<br>Date | HsOb reads/<br>total reads<sup>b</sup> |
| --- | --- | --- | --- | --- | --- | --- | --- | --- |
| H4 | 31°54'07.5"N<br>130°50'06.2"E | Hayashida | 68.8 | 3.2 | 2.1 | 1.3 | 10-Mar-2017 |  |
| H5 | 31°54'07.5"N<br>130°50'06.2"E | Hayashida | 69.7 | 3.1 | 2.0 | 1.8 | 10-Mar-2017 | 56/<br>1,734,416 |
| T1 | 31°54'37.7"N<br>130°49'00.6"E | Tearai | 92.1 | 2.9 | - | 0.0 | 09-Mar-2017 |  |
| T2 |  | Tearai | 95.9 | 2.1 | - | 0.0 | 09-Mar-2017 |  |
| T3 |  | Tearai | 94.4 | 2.4 | 0.0 | 0.0 | 09-Mar-2017 |  |
| T4 |  | Tearai | 92.8 | 2.7 | 0.0 | 0.0 | 09-Mar-2017 |  |
| Y66 | 31°55'03.8"N<br>130°48'40.4"E | Yunoike | 68.7 | 2.7 | 2.1 | 0.0 | 10-Mar-2017 |  |
| Y80 |  | Yunoike | 75-86<sup>a</sup> | 2.5 | 1.5 | 0.0 | 10-Mar-2017 |  |
| Y86 |  | Yunoike | 86.5 | 2.5 | 0.0 | 0.0 | 10-Mar-2017 |  |
| Oi | 32°44'25.3"N<br>130°15'48.4"E | Unzen | 79.3 | 2.2 | 0.0 | 0.4 | 18-Nov-2015 | 408/<br>316,160 |
| Ob | 32°43'33.0"N<br>130°12'24.7"E | Obama | 72.8 | 7.9 | 0.0 | 0.0 | 17-Nov-2015 |  |

<sup>\*</sup>Data other than read counts are reproduced from Urayama et al. (Urayama et al., 2024).
<sup>a</sup>There were temperature gradients in the pool site: surface layer 75.0 °C; bottom layer 81.6 °C, 80.3 °C, 85.9 °C; middle layer 81.0 °C.
<sup>b</sup>Mapping: Length fraction=0.9, Similarity fraction=0.9

### Obelisk-like elements from hot spring dsRNA metatranscriptomes

The single relatively large cccRNA and a related sequence were detected in the dsRNA metatranscriptome from two geographically distinct sampling stations, Oi and H5 (see below, Table 1). The Oi and H5 hot springs are characterized by high temperatures (69.7°C and 79.3°C) and low pH (3.1 and 2.2), respectively. The rRNA of potential host microbial communities from Oi and H5 were both dominated by thermophilic bacteria of the family *Hydrogenobaculaceae* in the phylum *Aquificota* (49.8 and 98.9%, respectively) (Urayama et al., 2024).

The circular sequence obtained from the acidic hot spring Oi (79.3°C, pH 2.2) consisted of 867 nucleotides with a GC content of 49.8% and did not show significant similarity to the sequences in the NCBI nucleotide (nr/nt) database. The secondary structure of the HsOb sequence predicted using RNAfold at the default temperature (37°C) resembled a long, unbranched rod with extensive intramolecular base pairing (68%) (Fig. 2A). At the physiologically-relevant temperature of 80°C, the predicted long rod-like structure was retained, although with a considerable decrease in base pairing. To further characterize the cccRNA, self-cleaving ribozymes were predicted using RNA sequence and secondary structure covariance models using Infernal (Nawrocki and Eddy, 2013); however, no known self-cleaving ribozymes were identified.

**Fig. 2:**
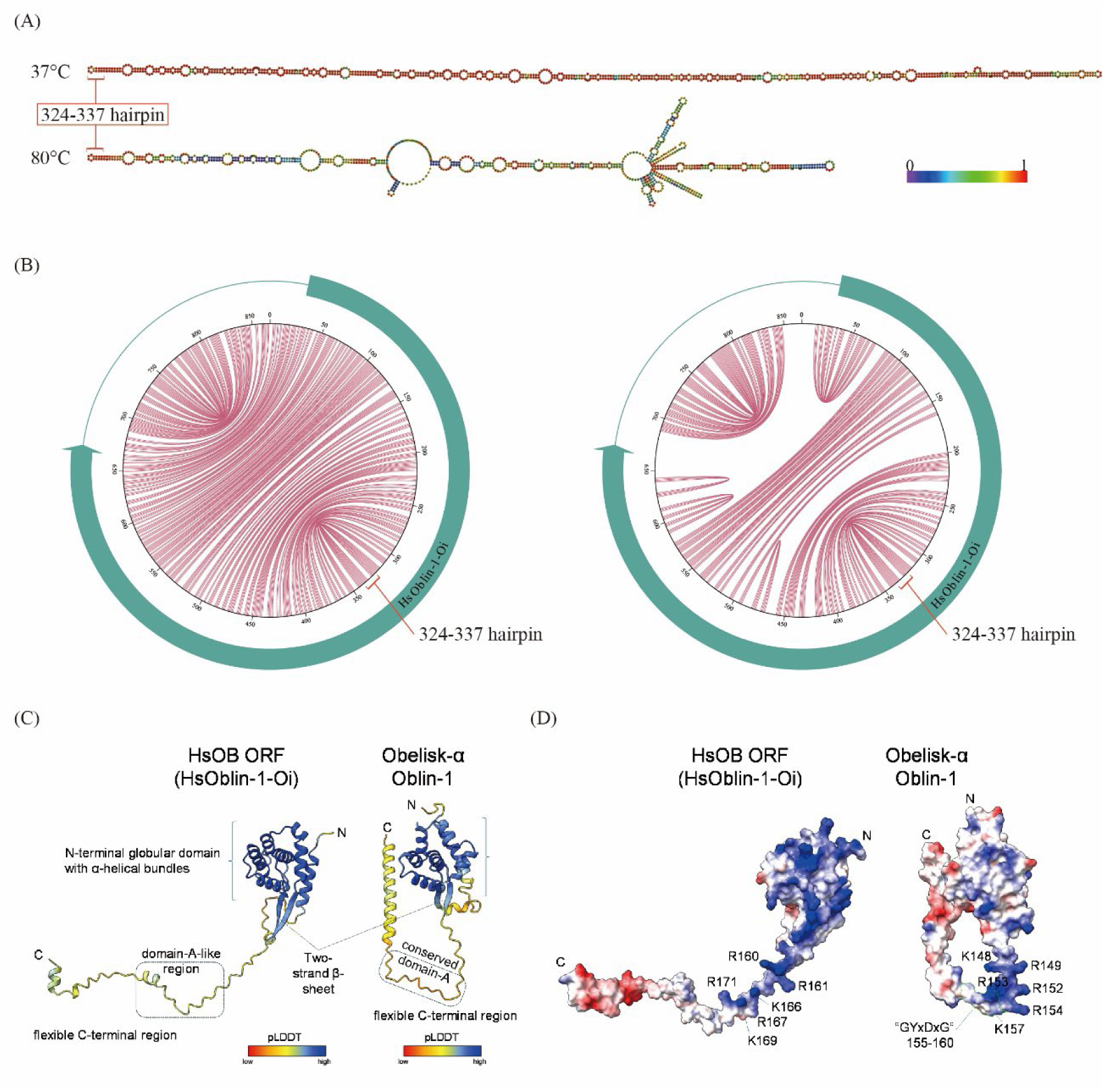
Characteristics of HsOb-Oi. (A) Schematic of the sense secondary structure derived from HsOb at 37°C and 80°C predicted by RNAfold. The ruler color indicates base-pairing probabilities. (B) Jupiter plot of HsOb at 37°C and 80°C and predicted open reading frame. (C) AlphaFold2 structural models of HsOblin-1-Oi and Oblin-1 from the Obelisk-α. Ribbons are colored by pLDDT scores and the color key is shown. (D) Electrostatic surface potential of the HsOblin-1-Oi and Oblin-1 protein models. Blue and red represent positive and negative changes, respectively. Positively charged residues around the domain-A region are labeled. The GYxDxG motif in Oblin-1 is highlighted by a green border line.

The cccRNA encompasses a single long ORF encoding a putative protein of 213 amino acids, which occupies approximately three-quarters of the nucleotide sequence (Fig. 2B). Although the amino acid sequence of the putative ORF product did not show significant similarity (BLASTP *E*-value ≤ 1 × 10^−05^) to the NCBI nr protein sequences, structural modeling using AlphaFold2 (AF2) and AlphaFold3 (AF3) yielded confident models of the HsOb-encoded protein (AF2, pLDDT=84.5, ptm=0.62; AF3, pLDDT=70.4, ptm=0.59; Fig. 2C and S1A). Comparison of the AF2 models of the HsOb-encoded protein and Oblin-1 revealed notable structural similarity (Root Mean Square Deviation of 3.817 Å as calculated using the MatchMaker tool of ChimeraX) despite the low sequence similarity (12.5% identity) (Fig. 2C). The shared predicted structural features included an N-terminal globular domain with α-helical bundles bookended by a two-stranded β-sheet, and an apparently flexible C-terminal region containing multiple positively-charged residues in the region around the position equivalent to the conserved domain-A in original Oblin-1, although there was no detectable amino acid sequence motif “GYxDxG” in this region of the ORF product (Fig. 2C-D, Fig. S4). In addition, we performed a FoldSeek search of protein structure databases using the confidently modeled N-terminal globular domain of cccRNA (aa51-148) as the query, but no significantly similar structures were detected in any of the databases (the best *E*-value was 0.31 for A0A481Z095; AlphaFold/UniProt50 v4, BFVD 2023_02, AlphaFold/Swiss-Prot v4, AlphaFold/Proteome v4, BFMD 20240623, Mgnify-ESM30 v1, PDB100 20240101, CATH50 4.3.0, GMGCL 2204; 3Di/AA, iterative search). Notably, however, these databases did not include the predicted structure of Oblin-1. These observations indicate that the cccRNA encoded a homolog of Oblin-1 with a high degree of structural conservation between the N-terminal alpha-helical bundle domains. Based on the shared genomic organization and the similarity of the encoded protein predicted structures, we concluded that the circular RNA from the Oi hot spring represents a distinct group of Obelisks (Zheludev et al., 2024) that we named Hot spring Obelisk-like RNA strain Oi (HsOb-Oi) (accession: XXX), denoting the encoded protein HsOblin-1-Oi.

During structural prediction optimization, we noted that adding RNA improved the confidence of HsOblin-1-Oi prediction by AF3. For example, supplying a generic 14-nt poly-U single-stranded RNA resulted in notably higher per-residue pLDDT scores, especially for the low-confident C-terminal region (Fig. S1B). The predicted aligned error (pAE) also showed substantial improvement in confidence with respect to the relative positions of the flexible C-terminal region in the presence of the RNA (Fig. S1C). To test a cognate RNA, the 14-nt terminal tetra-loop hairpin (324-337 of HsOb RNA) was selected based on the temperature-insensitive secondary structure (Fig. 2A-B). Similarly to the 14-nt poly-U RNA, the presence of the terminal hairpin RNA also led to the improvement both in pLDDT and pAE scores involving the C-terminal region of HsOblin-1, with a distinct RNA interaction surface predicted (Figs. S1B-C). In contrast, prebiously reported Obelisk-α Oblin-1 failed to generate a confident RNA-protein complex model either with a poly-U 14-nt RNA or a terminal hairpin RNA (Fig. S1D). These predictions suggest potential for multiple RNA-binding modes in HsOblin-1-Oi.

We then explored the sequence diversity of the HsOblin-1-Oi by searching the FLDS metatranscriptomes obtained from other hot springs. The amino acid sequence of the HsOblin-1-Oi was used as a query to search against the SPAdes-assembled contigs using BLASTX. This search yielded an additional HsOblin-1 sequence (*E*-value = 1 × 10^−108^; 71.7% pairwise amino acid sequence identity) encoded by a contig from sample H5. The H5 contig, denoted HsOb-H5, was incomplete (we could not recover circular sequence), but encoded a full-length HsOblin-1-Oi homolog which we denoted HsOblin-1-H5 (accession: XXX).

In an attempt to assign the host for HsObs, we searched their sequences against 919 CRISPR spacer sequences obtained by metagenomic DNA sequencing of the hot spring samples (Urayama et al., 2024) as well as 40,704 *Sulfolobales* spacers from the Beppu hot springs (Medvedeva et al., 2019). However, no spacers matching the HsOb genomes were identified.

### Major expansion of the Obelisk and Oblin-1 diversity

To identify additional Obelisks and Oblins related to HsOblin-1 and the prototype Oblin-1 (Zheludev et al., 2024), we mined ∼ 8.9 million putative cccRNAs from more than 5,000 metatranscriptomes from a recent study (Lee et al., 2023) and additional ∼4 million putative cccRNAs from about 2,000 assembled metatranscriptomes which more recently became available at IMG/MER (Chen et al., 2021), for homologs of Oblin-1 and HsOblin-1-like proteins using an iterative search procedure (see Methods). This search revealed 6,867 deduplicated Obelisk sequences encoding 5,009 deduplicated Oblin-1 and HsOblin-1 homologs. The retrieved homologous sequences were supplemented with about 1700 previously identified Oblin-1 centroid proteins (Zheludev et al., 2024), the FLDS identified HsOblin-1-Oi and -H5, and 5 (1 full-length and 4 partial sequences) HsOblin-1 proteins identified by BLASTX in Yellowstone hot spring sequencing projects (see Methods), bringing the total number of Oblin-1 homologs to 6,443 deduplicated sequences. The Obelisk set was clustered at 80 % average nucleotide identity (ANI) to follow the previously proposed nomenclature (Zheludev et al., 2024), resulting in 1774 clusters spanning known and newly identified Obelisks and 2060 clusters without known Obelisks, expanding the Obelisk diversity about two-fold at this level (see Methods).

This protein set was clustered by sequence similarity, yielding 111 clusters (see Methods). The relationships among these clusters remained uncertain because of the low sequence similarity approaching the limit of the HMM comparison resolution. The majority of large clusters (10 and more members) contained known Oblin-1 proteins, in addition to those discovered in this work, but 3 clusters consisted of new sequences only (Fig. 3, Clades C1-3), including the HsOblin-1-like cluster, C3, and one cluster contained only 4 known centroids (O18). The C3 cluster containing HsOblins and Yellowstone HsOblin-1 consisted of 44 sequences. For each cluster, a phylogenetic tree was built, midpoint rooted, and grafted onto the similarity dendrogram (Fig. 3A). Structure prediction from a representative set of diverse sequences across each clade supported the presence of an Oblin1-like fold in all these proteins although the two-stranded β-sheet was not always predicted with high confidence. Clustering 40 putative Oblin-1 sequences from a recent study (Lopez-Simon et al., 2025) with the present set at 50% sequence identity indicated that 19 of the 40 mapped to clade O13, 6 to O18 and 15 represented partial or diverged sequences that could not be mapped. Notably, for clade O18, a third beta-strand was predicted, interacting with the classical beta sheet clasp (Fig. 3B). Proteins in this clade are most closely related to a distinct set of several Oblin-1-like proteins reported by Zheludev and colleagues (Zheludev et al., 2024) which also shows a third beta-strand in their structure prediction (O16 and O17 in Fig. 3A), therefore substantially expanding this Oblin-1-like protein clade. Although structure comparison with Dali can yield low z-scores at the border of similarity detection even when comparing only the N-terminal globular domain with α-helical bundles bookended by the two-stranded β-sheet, the topology of the alpha-helical globular domain is conserved across the different newly discovered Oblin-1 cladess (Fig. S2). The majority of the cccRNAs encoding Oblin-1 homologs were within the 800-1,400 nt size range. The Oblin1-like ORF is typically between 400-1,000 nt long, resulting in a cccRNA coverage with the Oblin-1 ORF between 0.5-0.9 for the majority of sequences (Fig. 3C).

**Fig. 3:**
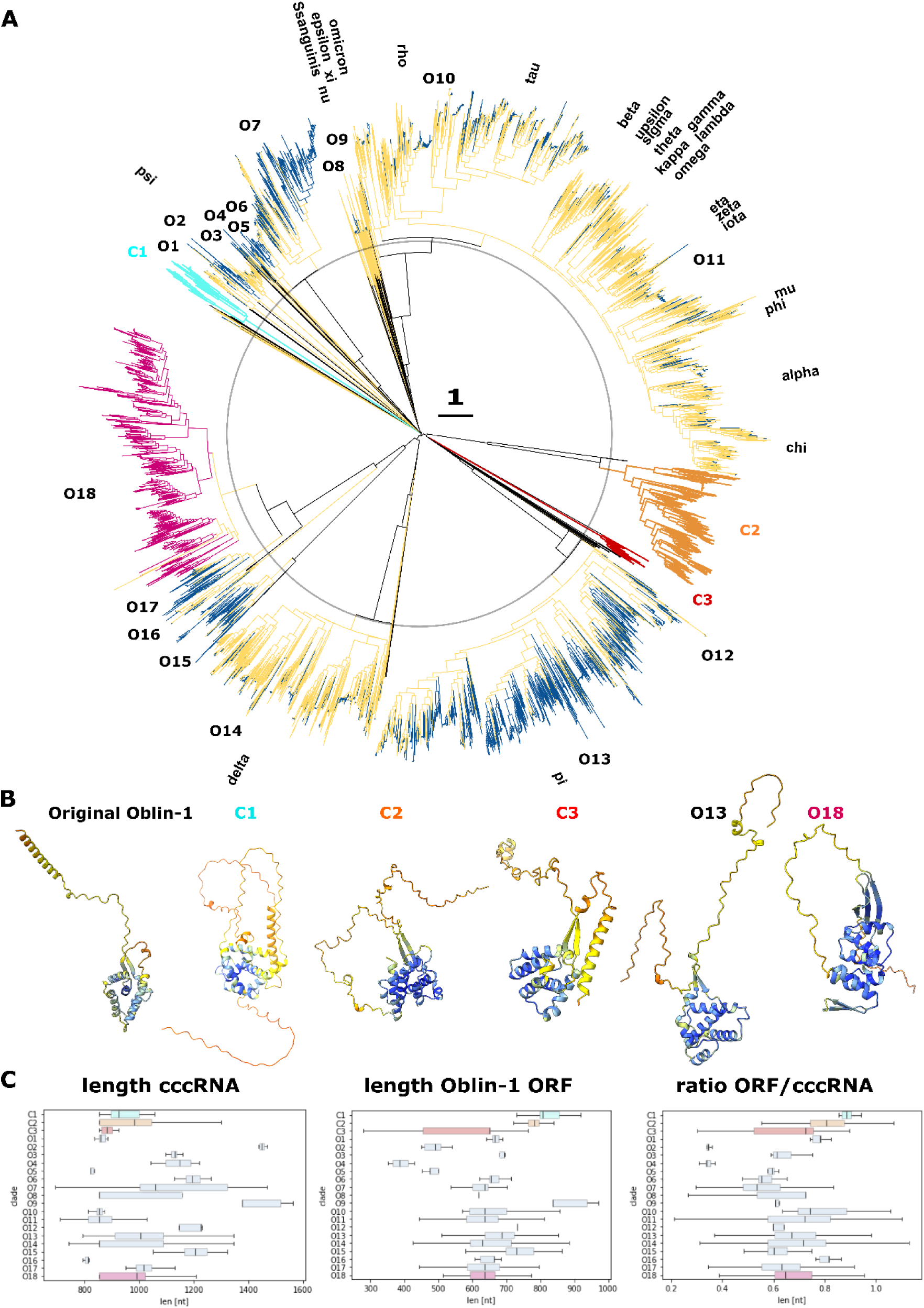
Overview of known and newly identified Oblin-1 proteins. (A) Inner part (till black circle): dendrogram based on pairwise hhsearch scores (including artificial low score in case no relationship was detected by hhsearch) of Oblin-1 clusters. Outer part: Midpoint routed FastTree phylogenetic trees based on alignment of Oblin-1 sequence which could be aligned confidently in an iterative procedure with hhalign. Yellow clades indicate those related to known Oblin-1 proteins (centroids of (Zheludev et al., 2024)) with dark blue terminal branches indicating sequences found in this study. Three larger clades (C1, C2 and C3) including HsOblin-1-like (red, C3) don’t harbor any known centroid Oblin-1 proteins. O18 contains only four known Oblin-1 sequences. (B) Representative structure predictions per selected clades of known and novel Oblin-1-like proteins. (C) cccRNA length, Oblin-1 ORF length and their ratio (from left to right) per clade (for clades with 10 or more members).

The phylogenetic tree indicated clustering of the sequences originating from different ecosystems (Fig. 4, Fig. S3). Aquatic and terrestrial environments dominate the respective clades and, for the C3 clade, sequences found in hot spring ecosystems cluster together on the tree. Three other clades included few Oblins from hot springs, all from bioprojects related to Yellowstone national park hot springs (C2, O11 and O13 with 2, 2 and 1 Oblin, respectively). Otherwise, Oblins from aquatic and terrestrial environments dominated clusters C1, C2, O18 and O13, a clade including known Oblin-1 proteins of the phi group but otherwise substantially expanded in this study. In an attempt to gain additional information on putative bacterial or archaeal hosts, we performed a CRISPR spacer search (see Methods) for all newly identified Obelisks. Only one Obelisk matched 29 nucleotides of a single 32 nt spacer from an RNA-targeting CRISPR system (Cas-III-B) of the bacterium, *Marinomonas mediterranea* (*Gammaproteobacteria*).

**Fig. 4:**
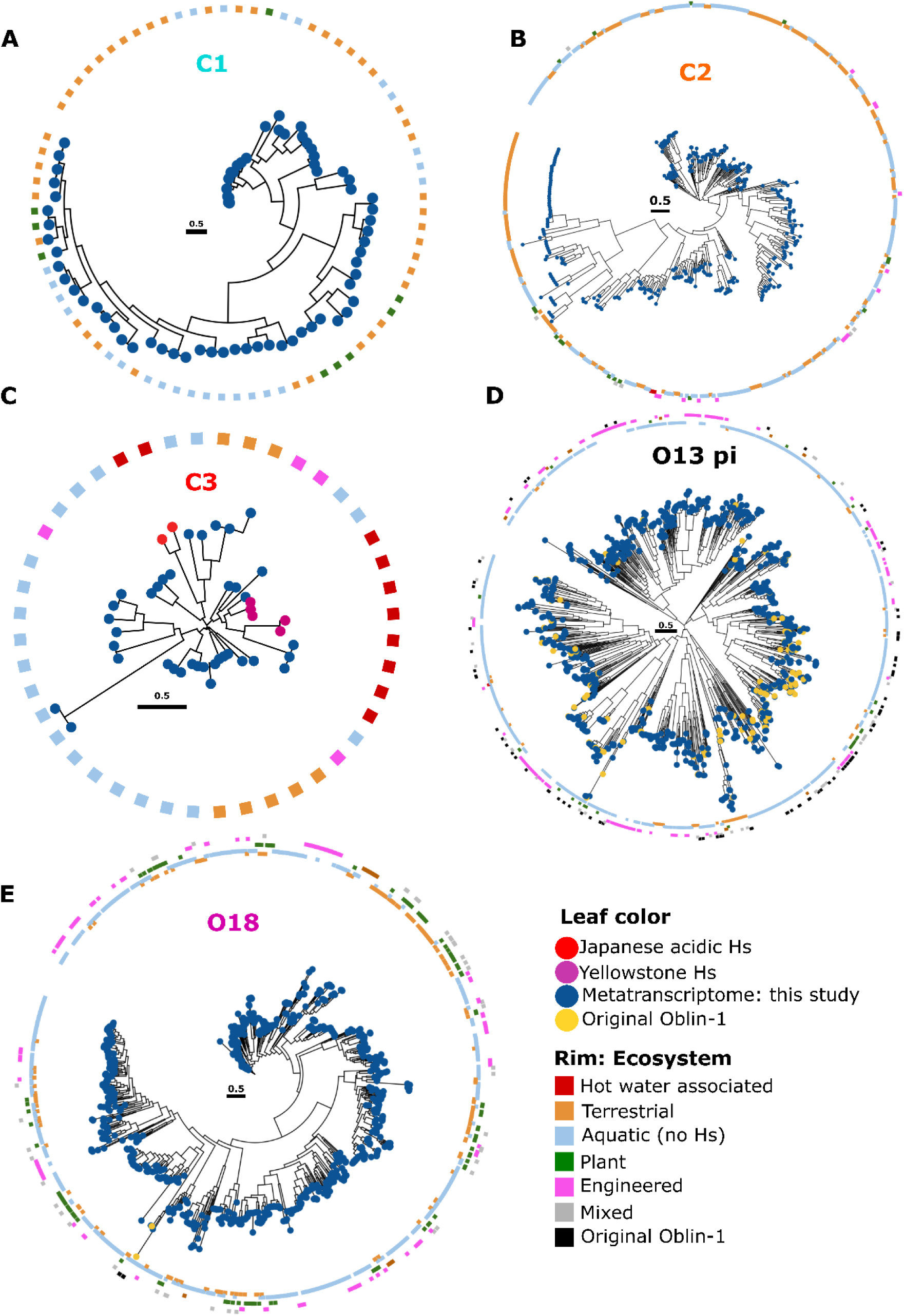
Ecosystems positive for Oblin-1 proteins from different subfamilies. (A-E) Unrooted phylogenetic Oblin-1 trees for selected clades are shown (clade nomenclature as in Fig. 3). Outer rim indicates the respective overarching ecosystem of the respective metatranscriptome study. Leaf color indicates Oblin-1 proteins discovered in this study (blue) or centroids from Zheludev et al. (Zheludev et al., 2024) (yellow).

Because HsOblin-1-Oi/H5 amino acid sequences appear to lack the “GYxDxG” motif which is the signature of domain-A in the original Oblin-1 proteins (Zheludev et al., 2024), we searched for sequence motifs conserved across the expanded cluster of 44 HsOblin-1-like proteins (C3) in the C-terminal region corresponding to the domain-A of Oblin-1s. HsOblin-1-like proteins encompassed three larger conserved regions that in the protein structure were located in positions corresponding to those of the conserved regions of known Oblin-1 proteins, two within the globular alpha-helical domain and one within the flexible C-terminal portion of the protein (Fig. 5). Similarly, no conserved GYxDxG motif could be detected for clades C1 and C2 as well as for O18, underpinning its status of a new Oblin-1 clade. Similar to HsOblin-1-like proteins, proteins from clades C1, C2 and O18 contain a stretch of variable length of positively charged residues. O18 contains in addition a highly conserved serine-glutamine pair, C1 contains two conserved serines (S(A/P/-)S) and C2 contains fewer positively charged residues but three tyrosines and two serines ((R/P)YSSY(C/T)KGYG) (Fig. S4). By contrast, Oblin-1 proteins from clades O16 and O17 retain the conserved GYxDxG motif.

**Fig. 5:**
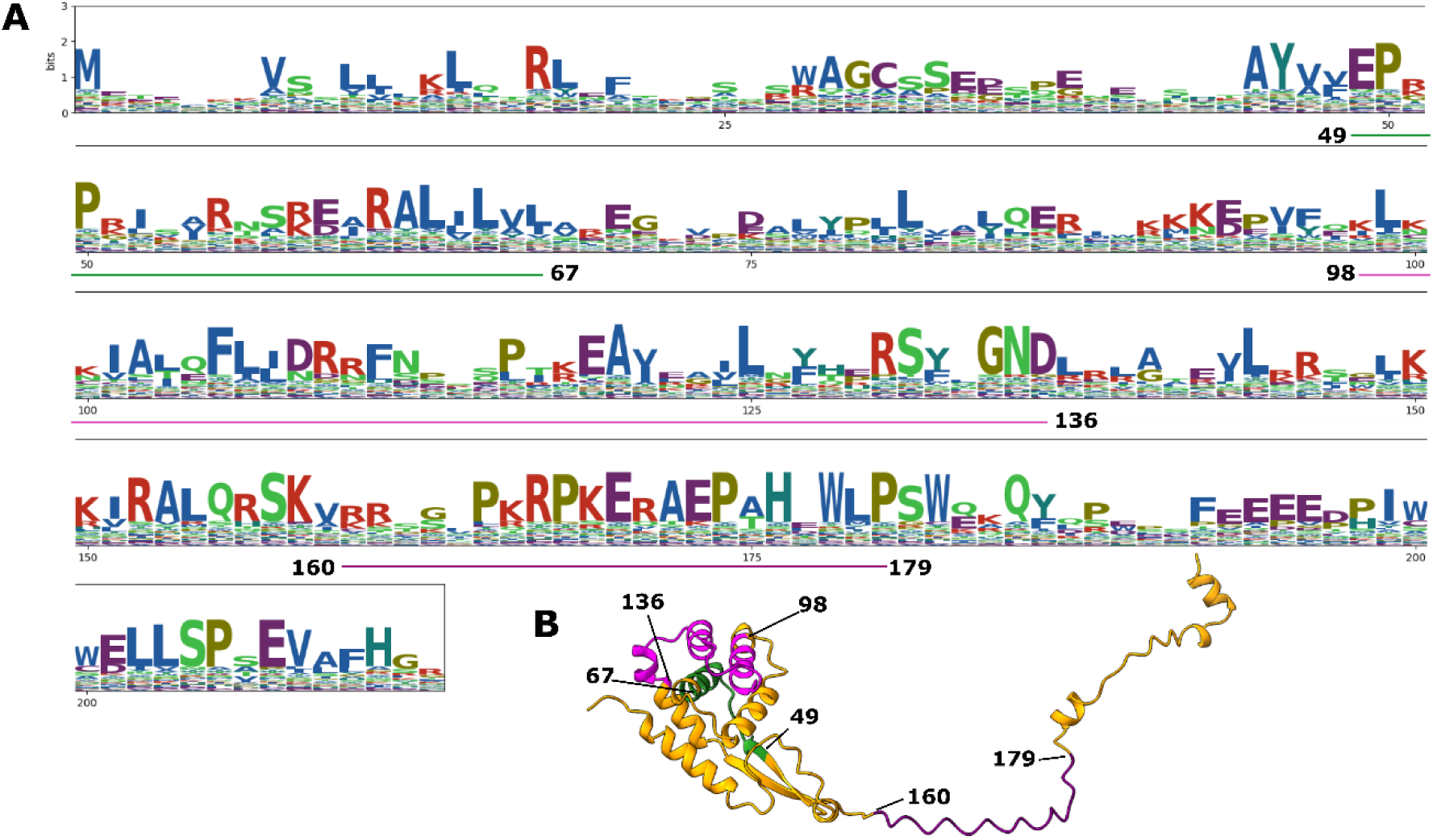
Conserved domains in HsObin-1 protein. (A) Amino acid logos (bits) obtained from the alignment of the 44 HsOblin-1-like proteins with conserved stretches underlined in green, pink and purple. Sites of the alignment matching HsOblin-1-Oi are shown. (B) AF2 HsOblin-1-Oi structure prediction with the respective conserved domains from (A) colored accordingly.

The secondary structures of Obelisk RNAs from hot springs and other newly identified clades were predicted using RNAfold (Lorenz et al., 2011). As expected, given the previously predicted rod-like structure of Obelisks as well as many other cccRNAs replicons, stable structures were predicted, the average minimum free energy for the majority of the clades being below −300 kcal/mole (Fig. S7A, B). In line with the predicted high stability of the Obelisk RNAs, the average fraction of paired bases was between 68-72 % except for clade O9 (64 % paired bases, sense strand) (Fig. S5C, D).

We searched all Obelisks for the presence of ribozymes using Infernal (Nawrocki and Eddy, 2013). Hammerhead ribozymes (HHR) were predicted in more than 800 Obelisks, mapping to more than 600 leaves in the Oblin-1 tree. The majority of these were HHR3 located in the strand complementary to the Oblin-1 encoding strand. Only a few HHRs other than HHR3 or ribozymes in the sense strand were predicted (Fig. S6). Mapping the ribozymes in relation to the Oblin-1 ORF indicated that the majority (70%) is on the antisense strand, non-overlapping with the Oblin-1 ORF and has a median distance of 51 nucleotides to the 5’ end of the Oblin-1 ORF. Other positions including completely nested ribozymes inside the Oblin-1 ORF were detected at lower frequencies (Fig. S7). The HHR3 were predicted mainly in Obelisks of the Oblin-1 clade O18 and associated clades O16 and O17, which is compatible with the original observations of Zheludev and colleagues (Zheludev et al., 2024). Also, the majority of ribozymes discovered here are found in comparable positions as reported by Zheludev and colleagues: a non-overlapping antisense HHR3 ribozyme in close proximity to the 5’ end of the Oblin-1 ORF and, if detected, a nested sense HHR3 ribozyme at the 3’ end of the Oblin-1 ORF.

### Diversity of non-Oblin-1 proteins encoded by Obelisks

The mean HsObs cccRNA size was 892 nt (range of 853-995 nt), whereas the mean HsOblin-like ORF length including the complete one from the Yellowstone hot spring metatranscriptome was 599 nt (range of 279 - 765 nt). ORFs shorter than 350 nt encode partial Oblins covering the C-terminus, indicating the possibility of alternative start codon use. The mean cccRNA coverage by the HsOblin-1-like ORF was 67% (range of 30-82%), leaving room for potential additional proteins. Comparable observations were made for cccRNAs from other Oblin-1 clades. To investigate the coding potential of Obelisks, amino acid sequences encoded by additional ORFs of the newly discovered and original Obelisks were extracted (see Methods). The 27,926 dereplicated putative non-Oblin-1 proteins were clustered by similarity (40% identity, 70% coverage), resulting in 18,609 clusters including 14,381 singletons. Of the 4,228 non-singleton clusters (2 to 48 members), 545 contained five or more members. The protein sequences from the non-singleton non-Ooblin-1 clusters were aligned, converted into HMM profiles and compared to each other using hhsearch. Cluster representatives were modeled using AlphaFold3 and compared to each other using foldseek. Structure comparison did not result in meaningful clusters due to the low prediction confidence for the majority of the sequences (Fig. S8) and the simplicity of the folds (mostly, small all helical domains with or without disordered regions). Therefore, only profile comparison was considered to define homologous protein groups. Members of these groups were mapped onto the Oblin-1 phylogeny to identify non-Oblin-1 proteins associated with the respective Oblin-1 proteins.

None of the putative non-Oblin-1 proteins encoded by HsObelisks or other new clades showed significant sequence similarity to the putative second protein encoded by the prototype Obelisks, Oblin-2. Furthermore, Oblin-2 proteins identified by PSI-BLAST using the prototype Oblin-2 as the query showed a patchy representation across the Oblin-1 tree and appeared mainly in subclades of Oblin-1 clades O11 and O14 (Fig. S9). In contrast, several other clusters of putative non-Oblin-1 proteins were strongly associated with distinct Oblin-1 branches. In particular, 10 non-Oblin-1 protein clusters mapped to Oblin-1 clades in which 70% or more of the Obelisks encode the respective non-Oblin-1 protein (Fig. S9 groups 0-9). Further, the two largest non-oblin-1 structure clusters mapped to Oblin-1 clades with a lower coverage (Fig. S9, lc1 and lc2). Inspection of the predicted structures of putative non-Oblin-1 proteins and the genome organizations of the respective Obelisks highlights 6 non-Oblin-1 protein clusters with confidently predicted structures and no overlaps with the Oblin-1 ORF, all associated with Oblin-1 clade O18 (Fig. S9 and S10 groups 0, 2, 6, 7 and lc2). In 5 of the 6 cases, the non-Oblin-1 ORF is encoded on the reverse strand compared to the Oblin-1 ORF (Fig. S10). Notably, in these 6 cases, the Oblin-1 and non-Oblin-1 ORFs are largely not base-paired in the cccRNA secondary structure prediction (both sense and antisense), demonstrating their capacity to evolve independently (Fig. S9B, S10).

Taken together, the above observations strongly suggest that at least the latter 6 clusters of non-Oblin-1 ORFs associated with O18 Oblin-1 clade actually encode proteins. Predicted folds of these non-Oblin-1 proteins are relatively simple alpha-helical structures including a beta-hairpin in three groups (Fig. S10, non-Oblin-1 groups 2, 7, and 9). Profile- and structure-based comparisons with various databases did not show confident hits for any non-Oblin-1 protein group. The actual existence of non-Oblin-1 Obelisk-encoded proteins apart from these 6 clusters, including Oblin-2, remains an open question.

## Discussion

Recent metatranscriptome mining has greatly expanded the known diversity of viroid-like cccRNAs (Lee et al., 2023; Lopez-Simon et al., 2025; Zheludev et al., 2024). To further explore these putative replicons in acidic thermal environments, we applied FLDS to samples from the hot springs in Japan. The HsOb was identified in a microbial community dominated by thermoacidophilic bacteria, and ribovirus RNA-dependent RNA polymerase (RdRP) sequences were also detected in the same samples (Urayama et al., 2024). Based on our results, the replication of HsOb is likely similar to that of previously reported Obelisks, although the involvement of RdRPs cannot be completely ruled out.

Previous metatranscriptome mining identified several groups of small, apparently non-coding cccRNAs in bacteria-dominated hot springs from Yellowstone, with the temperature of approximately 60°C (Koonin and Lee, 2025). For one of these non-coding cccRNAs, CRISPR targeting was demonstrated, clinching the case for a bacterial host (Koonin, 2024; Koonin and Lee, 2025). Here, we expand the diversity and upper temperature limit of Oblin-like cccRNA (79.3°C).

The discovery of HsOblin-1 combined with the extensive metatranscriptome mining for Oblins reported here nearly doubled members of this protein family, revealing a degree of diversity that extends beyond sequence similarity whereas the core protein fold is conserved, albeit with some variation, allowing recognition by structural comparison. Therefore, the observed 21 major clades (O1-18 and C1-3) can be considered provisional families within the Obelisk superfamily until a proper taxonomy is established. Notably, a long, disordered loop of Oblins showed clade-specific sequence conservation suggesting functional diversification. At least, for HsOblin-1, our structural modeling suggests binding of a specific element in the HsOb RNA, an interaction that might be important for HsOb replication. Also, other Oblin-1 clades without the canonical domain-A contained positively charged residues in this region, supporting their importance across Oblin-1 clades.

We did not detect broadly conserved Obelisk-encoded proteins other than Oblin-1. In particular, Oblin-2 predicted in the original Obelisk description was detected only sporadically in closely related Obelisks, casting doubt on the actual expression and functionality of this protein. However, in several new subfamilies of Obelisks, conserved small proteins with confidently predicted simple folds were identified. Thus, it appears that Obelisks as a class encode a single protein, Oblin-1, which is in all likelihood essential for their replication. However, some groups seem to have evolved additional protein-coding genes, possibly, encoding proteins involved in host-specific interactions.

The Oblin-1 homologs were detected in metatranscriptomes from a broad variety of environments suggesting that Obelisks replicate in diverse bacteria although specific hosts could not be readily identified apart from the original observation of Obelisk replication in *Streptococcus sanguinis* and the apparent association of HsObs with *Hydrogenobaculaceae* bacteria demonstrated here. Regardless, these findings, together with the recent demonstration of the high abundance of Obelisks in the oceans (Lopez-Simon et al., 2025), show that these are major components of the global microbiome that completely escaped detection prior to the recent advances of metatranscriptome mining.

The lifestyle(s) of the Obelisks and other cccRNA remains a major unresolved issue. Are these RNA plasmids or infectious agents, and to what extent do their replication strategies vary? These questions as well molecular mechanisms of Obelisk replication remain to be explored through identification of the hosts and laboratory cultivation.

## Methods

### FLDS analysis for plant leaves

Chrysanthemum Stunt Viroid (CSVd)-infected chrysanthemum leaves (0.2 g) were crushed using a mortar and pestle in the presence of liquid nitrogen, and total nucleic acids were extracted. After obtaining the dsRNA fraction using cellulose resin, residual non-dsRNA was degraded by treatment with DNase I and S1 nuclease. The purified dsRNA was fragmented using ultrasound (Covaris S220; Woburn, MA, USA), followed by ligation of the U2-primer to the 3’ end of the RNA. Reverse transcription was performed using the SMARTer system (Takara Bio, Kusatsu, Japan) with a primer (U2-comp-primer) containing the complementary sequence. After amplifying the cDNA by PCR, an Illumina sequencing library was prepared using the KAPA HyperPrep kit. Sequencing was carried out using Novaseq (Illumina) through an external company.

### Summary of dsRNA metatranscriptomes from geothermal hot springs in Japan

The hot spring dsRNA metatranscriptomes were constructed using the FLDS from high-temperature acidic hot spring microbiomes in Kyushu-island, Japan, as described previously (Table 1) (Urayama et al., 2024). The temperature and pH ranges for the samples ranged from 68.7 to 95.9°C and pH 2.2 to 3.7, respectively. Geochemical features and microbial community structures are also available. The genomes of RNA viruses were detected from the dsRNA metatranscriptomes dominated by thermophilic bacteria, whereas they were not from those dominated by hyperthermophilic archaea.

### Data processing

Trimmed reads were obtained using a custom Perl pipeline script (https://github.com/takakiy/FLDS) from dsRNA raw sequence reads (Hirai et al., 2021). The clean reads were subjected to *de novo* assembly using SPAdes (Bankevich et al., 2012) genome assembler v3.15.5 with metaplasmid mode. The sequences in the output file “before_rr.fasta” were used for ccfind program to find circular sequences with the following parameters: -L 130 -l 30. Sequences with significant similarity to known sequences were filtered out by BLASTn search against the NCBI nt database and BLASTx search against the nr database, using an *E*-value threshold of ≤ 1 × 10^−05^. In addition, sequences with low read coverage (<10 reads), short lengths (<200 nt), or insufficient secondary structure stability (base-pairing proportion ≤ 60% at 37°C, calculated using RNAfold (Gruber et al., 2008; Lorenz et al., 2011) were removed. To further select contigs with uniform and unbiased read distribution, we calculated two metrics: normalized coefficient of variation (CV) of read coverage and normalized entropy of read start positions. The normalized CV was calculated as the standard deviation divided by the mean of the read depth at each nucleotide position, and further normalized by the theoretical maximum CV expected for a sharp, single-peak coverage. This metric allows for fair comparison across sequences with different lengths and coverage depths. In contrast, normalized entropy was calculated based on the distribution of read start positions along each contig. Shannon entropy was first computed from the frequency distribution of start sites, then normalized by the maximum entropy possible for the given sequence length. This reflects the degree of randomness and diversity in where sequencing reads begin. Only contigs with a normalized CV < 0.6 and normalized entropy > 0.7 were retained for downstream analyses.

### Modeling protein structures for HsOblin-1-Oi

Initial structural predictions for HsOblin-1-Oi and Obelisk-alpha Oblin-1 were performed using ColabFold 1.5.1 installed locally through LocalColabFold (https://github.com/YoshitakaMo/localcolabfold) with default MSA setting and alphafold2_multimer_v3 as --model-type (Jumper et al., 2021; Mirdita et al., 2022). Number of recycles used for the resulting models of HsOblin-1-Oi (pLDDT=84.5, pTM=0.62) and Obelisk-alpha Oblin-1 (pLDDT=79.7, pTM=0.651) were 20 and 5, respectively. AlphaFold3 predictions for protein-alone and RNA-protein complex of HsOblin-1-Oi and Obelisk-alpha Oblin-1 were performed using the DeepMind’s AlphaFold server (https://alphafoldserver.com/, (Abramson et al., 2024). Model display, structural alignment, coloring, and figure preparation were performed using UCSF ChimeraX software (Pettersen et al., 2021).

### Screening public metatranscriptomes for HsOb-Oi/H5–related sequences

Metatranscriptomic datasets were mined from the NCBI Sequence Read Archive (SRA) by querying the taxonomic term “hot spring metagenome (txid433727)” together with the keyword “RNA”. The search, performed on 31 March 2025, returned 247 sequencing experiments. Datasets generated on non-Illumina platforms, those with insufficient read depth, and entries previously submitted by our group were discarded, leaving 154 samples corresponding to 197 sequencing runs (Table S2). For each run, raw reads were screened against the HsOb-Oi and HsOb-H5 reference sequences using BLASTN (BLAST+ v2.13.0, default parameters); no significant matches were detected. The open reading frames (ORFs) of HsOb-Oi and HsOb-H5 were then translated and used as database in BLASTX searches of the same read sets (*E*-value ≤ 1 × 10⁻⁵). This protein-level search identified eight samples (nine runs) that contained homologous sequences. Reads from these eight samples were assembled *de novo* in a sample-by-sample manner with MEGAHIT ver. 1.1.4, after quality filtering using Trimmomatic ver. 0.35, as detailed in a previous study (Nishimura and Yoshizawa, 2022). Contigs of each assembly were re-screened with BLASTX using the same *E*-value threshold. Five contigs showing significant similarity (*E*-value ≤ 1 × 10⁻⁵) to the HsOb-Oi/H5 ORFs were extracted. Of all the sequences, only one was successfully recovered as a circular sequence using ccfind and the sequence was circularly permuted not to cut off an ORF in the middle. Notably, all five contigs originated from samples of Yellowstone hot springs.

### Search for Obelisks and Oblin-1 homologs in the putative cccRNA dataset

To identify additional Oblins and Obelisks related to HsOblin-1 and classical Oblin-1, about 4.8 putative proteins (at least 60 aa long) from 8.9 million recently published (Lee et al., 2023) putative cccRNAs from more than 5000 metatranscriptomes were searched for the presence of HsOblin-1 and Oblin-1 homologs using an iterative search procedure. One round of PSI-BLAST (Altschul et al., 1997) was run with the two aligned HsOblin-1 sequences as the query, the significant matches (*E*-value ≤ 1 × 10^−05^) were aligned with MUSCLE5 (Edgar, 2022), and a HMMER profile build. The same procedure was followed for a prototype Oblin-1 (Obelisk_000001_000001_000035 from Zheludev et al.) to fetch homologous Oblin-1 proteins and the respective cccRNAs. The resulting HsOblin-1 and Oblin-1 HMMER profiles were enriched by a HMMER hmmsearch run (version 3.4, http://hmmer.org/) against the 4.8 million proteins derived from putative cccRNAs and against 1,700 centromer Oblin-1 proteins recently published (Lee et al., 2023; Zheludev et al., 2024). All confident HMMER hits (*E*-value ≤ 1 × 10^−10^) were kept, aligned with MUSCLE5 and final HMMER profiles build. For HsOblin-1, all 31 sequences stem from metatranscriptomics. For Oblin-1, 456 sequences stem from metatranscriptomics and 1,181 from centroids and three individual alignments and profiles were constructed using either only Oblin-1 sequences from metatranscriptomics or centroids or all combined. All final profiles were run again against the 4.8 million proteins from metatranscriptomics-derived cccRNAs (HMMER hmmsearch, default parameters). All about 2,600 HMMER hits for HsOblin-1-like and Oblin-1 were collected independent of their *E*-value, to allow discovery and inclusion of distantly related Oblin-1 proteins. This assemblage was enriched with about 1,700 Oblin-1 centroid proteins identified by Zheludev et al. (Zheludev et al., 2024), the two HsOblin-1 proteins identified by FLDS and five (1 full-length and 4 partial sequences) HsOblin-1 homologs identified in Yellowstone hot spring sequencing projects (see above). Proteins were subsequentially deduplicated (100% identity), leading to 3,454 deduplicated (Hs)Oblin-1 homologs.

This protein sequence set was analyzed to identify sequences which could be used as seeds for additional Oblin1-like profiles as well as to identify false-positives among the hits with high *E*-values. To these ends, proteins were clustered using MMSEQS2 (Steinegger and Söding, 2017) (0.5 sequence identity, 0.7 coverage, resulting in 1211 clusters), the sequences in each cluster were aligned using MUSCLE5 (Edgar, 2022) and an HMM profile was built for each alignment using HH-suite3 (Steinegger et al., 2019). In addition, protein structures were predicted for 1,211 representative sequences from the initial MMSEQS2 clusters using AlphaFold3 (see below). Clusters were compared to each other using HHSEARCH (Söding, 2005) and most related clusters aligned to each other by HHALIGN (Söding, 2005) as previously described (Wolf et al., 2018). In brief, sites in each cluster alignment containing more than 67% gaps were temporarily removed and pairwise scores obtained from an all-vs-all comparison with HHSEARCH. Pairwise scores d_A,B_ of cluster A and B were converted into distances using the formula d_A,B_ = −log[S_A,B_/min(S_A,A_,S_B,B_) in which S_A,B_ is the HHSEARCH score between the respective pair and S_A,A_ and S_B,B_ the self-score. Matrix of pairwise scores was used to construct a UPGMA dendrogram using the R package hclust with the argument ‘method’ set to “average” (=UPGMA). Clusters corresponding to related, shallow tips of the dendrogram were iteratively aligned to each other using HHALIGN. Further, neighboring clusters were only aligned if i) the pairwise distance did not exceed 2.2 or ii) the alignment coverage was at least 0.66. The procedure was iterated for 7 cycles, after which merging of alignments decreased the alignment quality (visual inspection). This procedure resulted in 153 clusters of aligned sequences of which 140 contained less than 10 sequences (spanning 261/ 3,454 sequences). Aligned clusters were converted into HMMER profiles which were used as queries to search against the complete set of cccRNA-encoded proteins to identify clusters that could be used to identify additional, distantly related Oblin-1 homologs. Sequences in clusters that initially contained fewer than 10 sequences were re-aligned with confident hits (*E*-value ≤ 1 × 10^−08^), HMMER HMMs were build and re-run against the full cccRNA-encoded protein set. All additional hits not present yet in the initial assemblage with an *E*-value below 1 × 10^−08^ were kept.

Structure comparison as well as cluster-specific pairwise HMM comparison indicated that 75/153 aligned clusters with 10 or less sequences were likely false positives and were accordingly discarded (86 sequences across the 75 clusters, mainly, singletons). Additional distant Oblin1-like sequences not covered by the first original HsOblin-1 and Onlin-1 profiles were detected for 32 of the aligned clusters. Overall, the enrichment runs resulted in additional 1,750 deduplicated Oblin1-like proteins and five clusters, including one spanning HsOblin-1 proteins, not containing known Oblin-1 centroid proteins. These five clusters were aligned and additional HMMER profiles built. To find additional Obelisks, over 2,000 assembled metatranscriptomics published after the study of Lee et al. in 2023 (Lee et al., 2023) were searched for cccRNAs (see below, about 4.1 million unique cccNAs), ORFs extracted and 2.2 million proteins of at least 60 aa searched with the different Oblin-1 HMMER profiles This resulted in 1,467 additional unique (Hs)Oblin1-like proteins.

To analyze the overall relationships across all identified 5,009 unique (Hs)Oblin-1 homologs the FLDS identified HsOblin-1-Oi and -H5, the five Oblin-1 proteins identified in a Yellowstone hot spring metatranscriptome and the 1,700 known centroid Oblin-1 proteins, MMSEQS2 clustering and the above described iterative pairwise alignment of clusters using HHSEARCH and HHALIGN were repeated. Finally, 111 aligned clusters were obtained. The phylogeny for each cluster was constructed using FastTree (Price et al., 2009, 2010) (-wag -gamma options)-after removing sites with more than 90% gaps from the respective MSAs. The obtained trees were midpoint rooted and grafted to the HHSEARCH based UPGMA dendrogram of the aligned clusters. Protein structures were predicted for a representative set of diverse sequences across each clade using AlphaFold3 (see below for details).

### Tree visualization

Phylogenetic trees and dendrograms were visualized with iTol web interface (https://itol.embl.de/) (Letunic and Bork, 2024).

### Search for additional cccRNAs in metatranscripomes

Assembled metatranscriptomics deposited after the publication of Lee et al. (Lee et al., 2023) were retrieved from IMG/MER (Chen et al., 2021) and putative cccRNAs retrieved with vdsearch (Lee et al., 2023)(https://github.com/Benjamin-Lee/vdsearch). Monomeric cccRNAs were further dereplicated with circKit (https://github.com/Benjamin-Lee/circkit).

### Prediction of protein-coding genes

ORFs on cccRNAs were identified predicted in all 6 frames by circKit (https://github.com/Benjamin-Lee/circkit) default settings (start-stop, accounting for circularity and stopping after three wrappings of the genome in case no stop is present). For the Oblin-1 search, only putative proteins of at least 60 aa were considered. For non-oblin-1 proteins, all proteins of at least 40 aa from Obelisks encoding a (Hs)Oblin1-like protein were considered.

### Non-Oblin-1 protein analysis

Putative proteins of at least 40 aa encoded on Obelisks besides Oblin-1 (named here non-Oblin-1 proteins) were retrieved from all six frames and clustered with MMSEQS2 (Steinegger and Söding, 2017) (default parameters except: min-seq-id 0.4 -c 0.7). For clusters with at least two members, protein structures were predicted with AlphaFold3 (see below). Structures were clustered with foldseek (Van Kempen et al., 2024) (coverage -c 0.8). Structure-based clusters were inspected manually, indicating that clustering is mainly based on small, simple secondary folds such as single alpha-helices. Therefore, to avoid clustering of unrelated proteins, only sequence-based clustering was considered further. Sequence-based clusters with more than 5 members were aligned with MUSCLE5 (Edgar, 2022) and compared with HHBLITS (two iterations) against the following broad HHSUITE databases: pdb70 (Berman et al., 2000), pfama (Mistry et al., 2021), scope (Chandonia et al., 2022; Fox et al., 2014), ECOD (Cheng et al., 2014); and two HHSUITE databases for RNA virus proteins: nvpc (Neri et al., 2022) and viral (Wolf et al., 2020). Only 4/18,609 clusters showed hits with a probability higher than 90 and these 4 hits were mainly mapping back to short helices in the non-Oblin-1 proteins likely unrelated to the function associated with the hits. Further, all putative non-Oblin-1 proteins (independent of the sequence-based clustering) were searched against the HMMER pfam database (based on pfam34.0), retrieving no conclusive hits.

Sequence-based clusters containing at least 20 proteins were projected on the Oblin-1 based tree. The deepest node being parental to all Obelisks encoding one non-Oblin-1 protein of the respective cluster was identified and the fraction of leaves encoding for a cluster-specific non-Oblin-1 proteins in this clade retrieved. Further, the mean pairwise distance of all leaves in this clade was retrieved. We excluded clades with a mean pairwise distance of less than 0.1 to avoid the detection of false-positive non-Oblin-1 ORFs by the high similarity of the underlying Obelisk sequences. Further, clades with a coverage of at least 70% of leaves encoding for the putative non-Oblin-1 proteins were inspected in detail. In addition, the two clades with more than 100 putative non-Oblin-1 proteins were also inspected, although providing a lower coverage. This retrieved 12 sets of putative non-Oblin-1 proteins which are evolutionary related to the respective Oblin-1 protein. Protein structures and ORF positions were visually inspected for these 12 clades. Further, the percentage of predicted base-pairing between the non-Oblin-1 and the Oblin-1 ORF in the secondary structure prediction (both sense and antisense) was extracted from the RNAfold-based prediction (see below).

In addition, the full non-Oblin-1 protein set was searched for Oblin-2 proteins described previously by Zheludev and colleagues. Therefore, a prototype Oblin-2 sequence (‘MDS VQI LRK KIL KNE EQR EFL LKK IGN LEY EIN NLE HKI ENQ QRV LQN LLR EK’) (Zheludev et al., 2024) was used as PSI-BLAST query and run until convergence (default settings except: -evalue 0.01 -outfmt 6 -max_target_seqs 50000 -num_iterations 0). Putative Oblin-2 (*E*-value ≤ 1 × 10^−03^) were mapped on the Oblin-1 phylogeny.

### Structure prediction, comparison and visualization for the expanded Oblin1-like protein family

Protein structures for the expanded (Hs)Oblin1-like proteins and non-oblin-proteins were predicted with a local version of AlphaFold3 (Abramson et al., 2024). Besides default settings, all dereplicated putative proteins from 8.9 million cccRNAs found in metatranscripomes (Lee et al., 2023) (majority are non-Obelisks) have been visible to AlphaFold3 during the MSA construction phase by concatenating them to the uniref90 file used by AlphaFold3 (uniref90_2022_05.fa). Chimera X (Pettersen et al., 2021) was used to visualize structures.

Selected (Hs)Oblin-1-like core structures (globular fold surrounded by the conserved beta sheets) have been compared pairwise (Fig. S2) using the Dali web server (Holm, 2022). The pairwise Dali translation-rotation matrix coordinates were used in Chimera X to superimpose structures using the ‘view matrix mod’ command.

Foldseek easy-search (Van Kempen et al., 2024) was used to search non-Oblin-1 proteins against several foldseek databases (pdb, alphafold2 proteome, alphafold2 swissport and big fantastic database). Foldseek search (-c 0.8) and clust (default) commands were used to compare non-Oblin-1 proteins against each other.

### RNA secondary structure prediction for Obelisks from metatranscriptomic data

RNA secondary structures of metatranscriptomic derived Obelisks and centroids were predicted with RNAfold implemented in the viennaRNA package (Lorenz et al., 2011), accounting for circularity (RNAfold-c) and predicting structures for both the sense and the antisense strand.

### cccRNA genome visualization and jupyter plots

Obelisk maps were plotted using the python library Plasmidviewer (https://github.com/ponnhide/plasmidviewer) which takes the coordinates from a GenBank formatted file. Obelisks were oriented in a way that the Oblin-1 ORF is always visualized on the sense strand. Jupyter plots of RNA secondary structure base pairs were plotted using the python script ‘jupiter’ (https://github.com/rcedgar/jupiter) which takes the base pairing coordinates from the RNAfold RNA secondary structure prediction.

### Ribozyme prediction

To detect ribozymes among the Obelisks detected in metatranscriptomic data, infernal (http://eddylab.org/infernal/ and (Nawrocki and Eddy, 2013)) as implemented in vdsearch (Lee et al., 2023). Running vdsearch infernal, default settings with a static ribozyme collection from ViroidDB (Lee et al., 2022).

### CRISPR spacer search

2x concatenated Obelisk sequences (both sense and antisense) have been used to search a local CRISPR spacer database for which all CRISPR Arrays in all bacterial and archaeal gemomic sequences in RefSeq as of May 2023 were predicted using minced (https://github.com/ctSkennerton/minced). BLASTN was used (blastn -query ip_fasta -task blatn -db sp2305.db -out op_file -evalue 1000 -word_size 4 -outfmt ‘6 qaccver saccver pident length mismatch gapopen qstart qend sstart send evalue bitscore nident slen’. Spacers covered by at least 90 % and at least 16 matching nucleotides have been inspected further.

### Visualization of sequence conservation

Visualization of sequence conservation of aligned Oblin-1 clusters as stacked sequence logos (information content as bits (Tareen and Kinney, 2020)) was done with the python package logomaker (Tareen and Kinney, 2020).

### Obelisk clustering

To follow the nomenclature proposed by Zehludev and colleagues, we clustered all known Obelisks (∼7000 from Zheludev et al. (Zheludev et al., 2024) and 40 from López-Simón et al. (Lopez-Simon et al., 2025) with the ones discovered here at 80 % nucleotide sequence identity (ANI) with circuclust (https://github.com/rcedgar/circuclust). Cluster members were then individually clustered at 95 % ANI to follow the nomenclature: Obelisk_X_Y_Z”, “X” denotes no. of cluster at 80% nucleotide identity, “Y” at 95 %,and “Z” as the identifier for all strains within a given species.

## Data availability

Datasets obtained in this study have been available in the Short Read Archive database (Accession No. DRA016131). Data including Obelisks and their proteins identified in this study is made available via zenodo (xxxxxx).

## Code availability

A custom code used in this study was made available in a git repository publicly available on GitHub at https://github.com/takakiy/FLDS (Cleanup_FLDS.pl) and https://github.com/yosuken/ccfind (ccfind).

## Acknowledgments

We gratefully acknowledge NARO GenBank for providing viroid strains used in this study. We thank Hiroko Mizukoshi and Tamaki Ichiki-Uehara for their experimental and technical support. This study was supported by Grants-in-Aid for Scientific Research on Innovative Areas from the Ministry of Education, Culture, Science, Sports, and Technology (MEXT) of Japan (Grant Nos. 25K22486 [for S.U., Y.M. and Y.N.], 23K18146 [for S.U. and Y.M.], 24K02083 [for S.U.], and 20K20377 [for T.N.]). This research was also supported in part by Lilly Endowment, Inc., through its support for the Indiana University Pervasive Technology Institute which provided supercomputing resources for protein structure modeling. This work utilized the computational resources of the NIH HPC Biowulf cluster (https://hpc.nih.gov). We appreciate Igor Tolstoy’s (NCBI, NIH) help in the construction of the CRISPR spacer database used in this work. We thank Benjamin Lee for his help with vdsearch and circkit. P.M. and E.V.K. are supported by the Intramural Research Program of the National Institutes of Health (NIH). The contributions of the NIH authors are considered Works of the United States Government. The findings and conclusions presented in this paper are those of the authors and do not necessarily reflect the views of the NIH or the U.S. Department of Health and Human Services.

## Author Contributions

All authors had a substantial contribution to this work. S.U., A.F., P.M., M.K., E.V.K. and T.N. were responsible for the design of the work, the acquisition, analysis and interpretation of data, drafted the initial work. S.U. and Y.M. performed experiments, and analyzed and interpreted the data. S.U., A.F., P.M., M.K., E.V.K. and T.N. substantively revised the work. S.U., A.F., P.M., Y.N., Y.T., S.M. and M.K. performed bioinformatic analysis. S.U., A.F., P.M., Y.M., Y.T., Y.N., S.M., M.K., E.V.K. and T.N. wrote the manuscript.

---

**Table S1:** Computed metrics reflecting the uniformity of read mapping across each candidate.

| Sample | Candidates | Normalized<br>CV<br>($< 0.6$) | Normalized<br>entropy<br>($> 0.7$) | Comment |
| --- | --- | --- | --- | --- |
| Leaves | **NODE_115** | **0.581** | **0.899** | CSVd |
|  | **NODE_392** | **0.327** | **0.930** | CSVd |
| Hot Spring | can01 | 1.140 | 0.483 |  |
|  | can02 | **0.564** | 0.631 |  |
|  | can03 | 1.629 | 0.451 |  |
|  | can06a | 1.532 | 0.634 |  |
|  | can06b | 1.370 | 0.629 |  |
|  | can07 | 0.755 | 0.457 |  |
|  | can09a | 1.487 | 0.667 |  |
|  | can09b | 1.013 | **0.751** |  |
|  | can11 | 0.693 | **0.711** |  |
|  | **can13** | **0.381** | **0.766** | HsOb |

**Table S2: 197 SRA sequencing runs used for sequence search.**

**Fig. S1:**
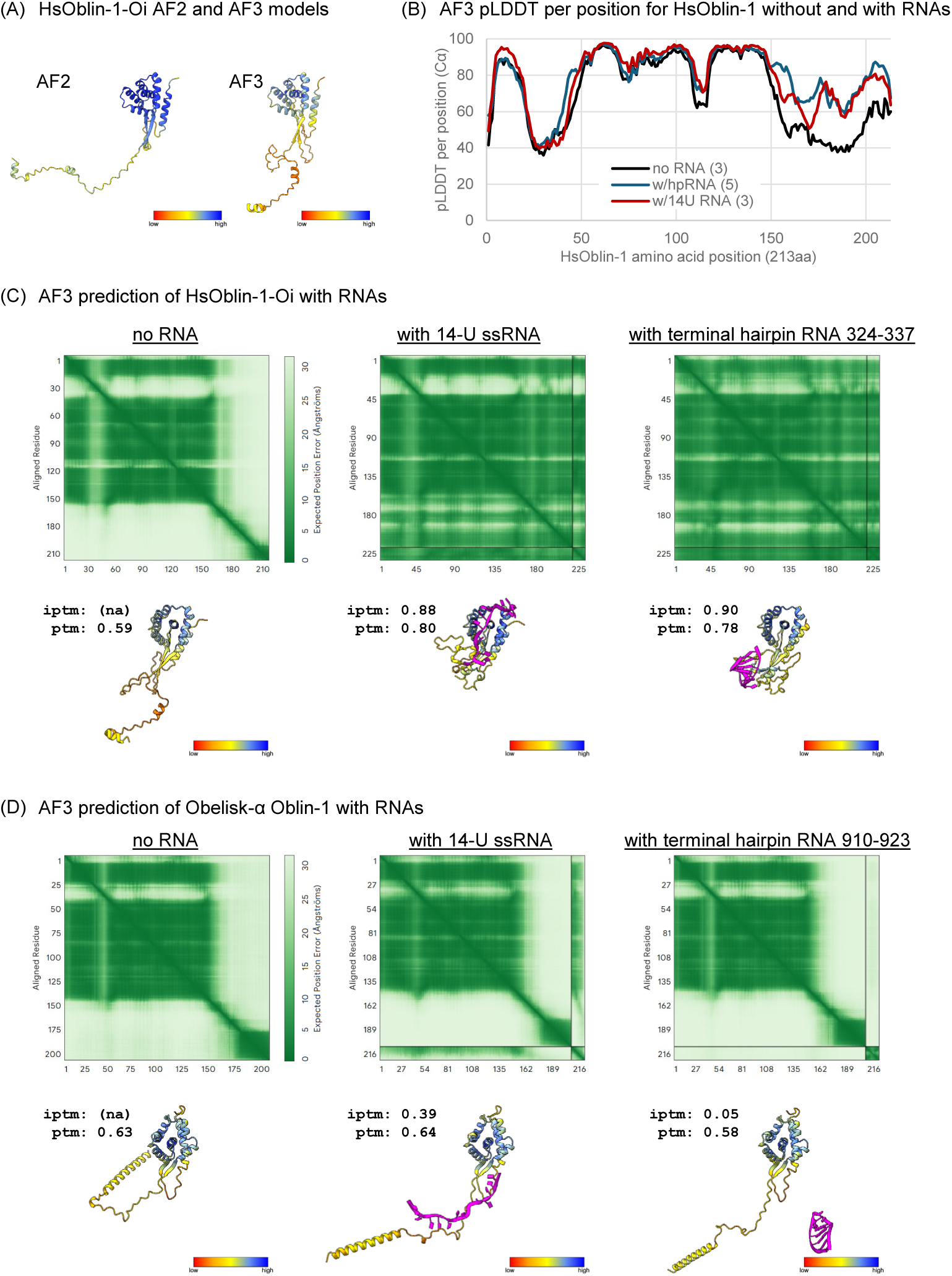
Improvement of AlphaFold3 HsOblin-1 structural modeling with RNAs. (A) HsOblin-1-Oi structural models by AlphaFold2 (AF2) and AlphaFold3 (AF3). (B) Summary of AF3 per-residue pLDDT scores for Cα atoms from HsOblin-1-Oi predictions without and with RNA. Numbers in parentheses show technical replicates with random seeds used to average the pLDDT score. (C-D) pAE plot and predicted models of representative AF3 predictions for HsOblin-1-Oi (C) and Obelisk-α Oblin-1 (D) without and with RNAs. The protein ribbons are colored by pLDDT. RNA chains are colored purple.

**Fig. S2:**
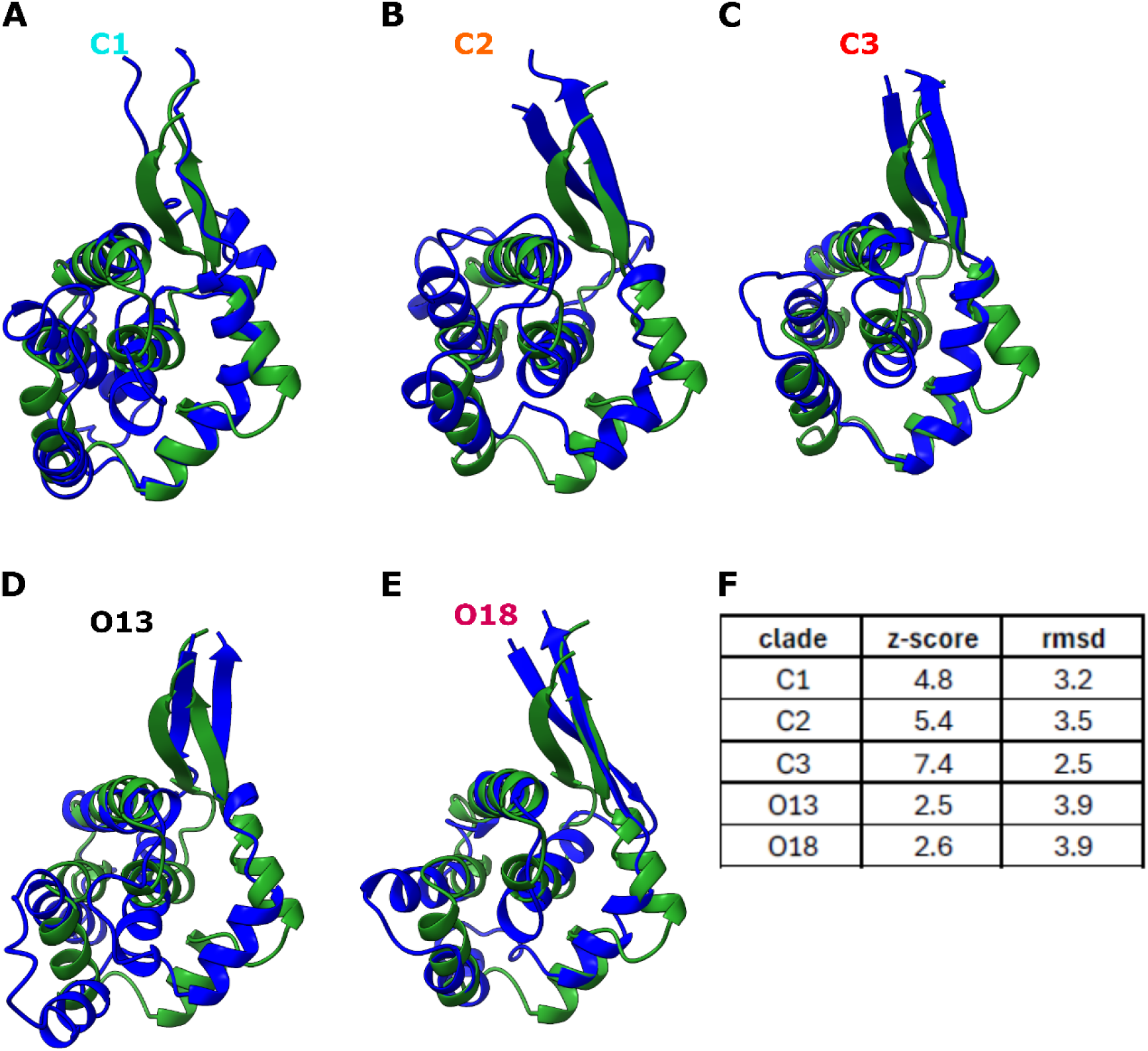
Structure comparison of different Oblin-1 subfamilies. (A-E) A representative Oblin-1 core structure (blue, globular fold flanked by the two beta-strands) of the indicated clade is superimposed to a predicted core structure of a classical Oblin-1 protein (green, globular fold flanked by the two beta-strands). (F) DALI Z-score and rmsd values of the pairwise compared core structures shown in A-E.

**Fig. S3:**
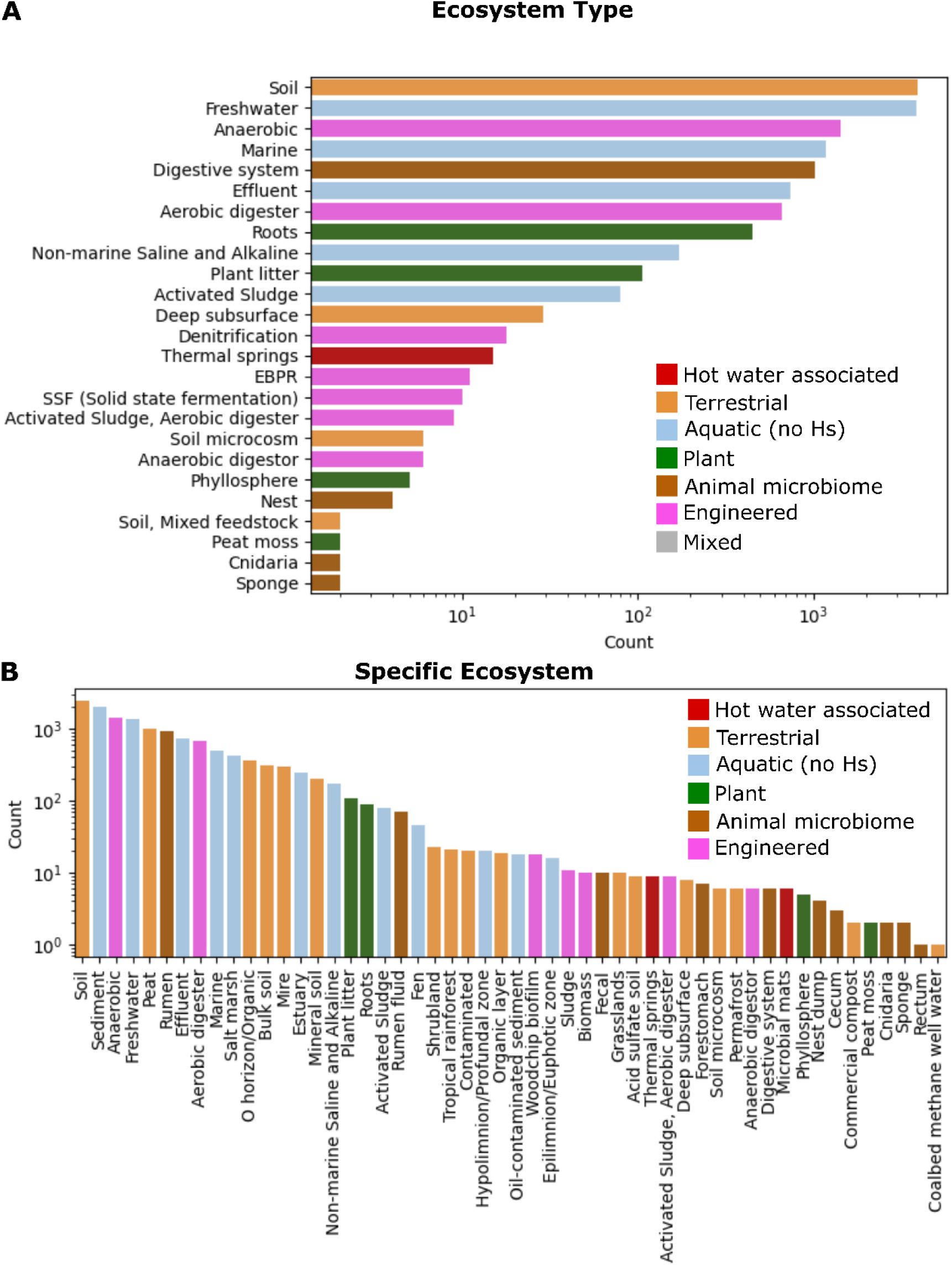
Ecosystem positive for Obelisks. Ecosystems in which Oblins from all major clades have been found in. Number of metatranscriptomes with Oblins by IMG metatranscriptome annotation for ‘ecosystem type’ (A) and ‘specific ecosystem (B). Bars are colored by overarching ecosystem complementary to Fig. 4.

**Fig. S4:**
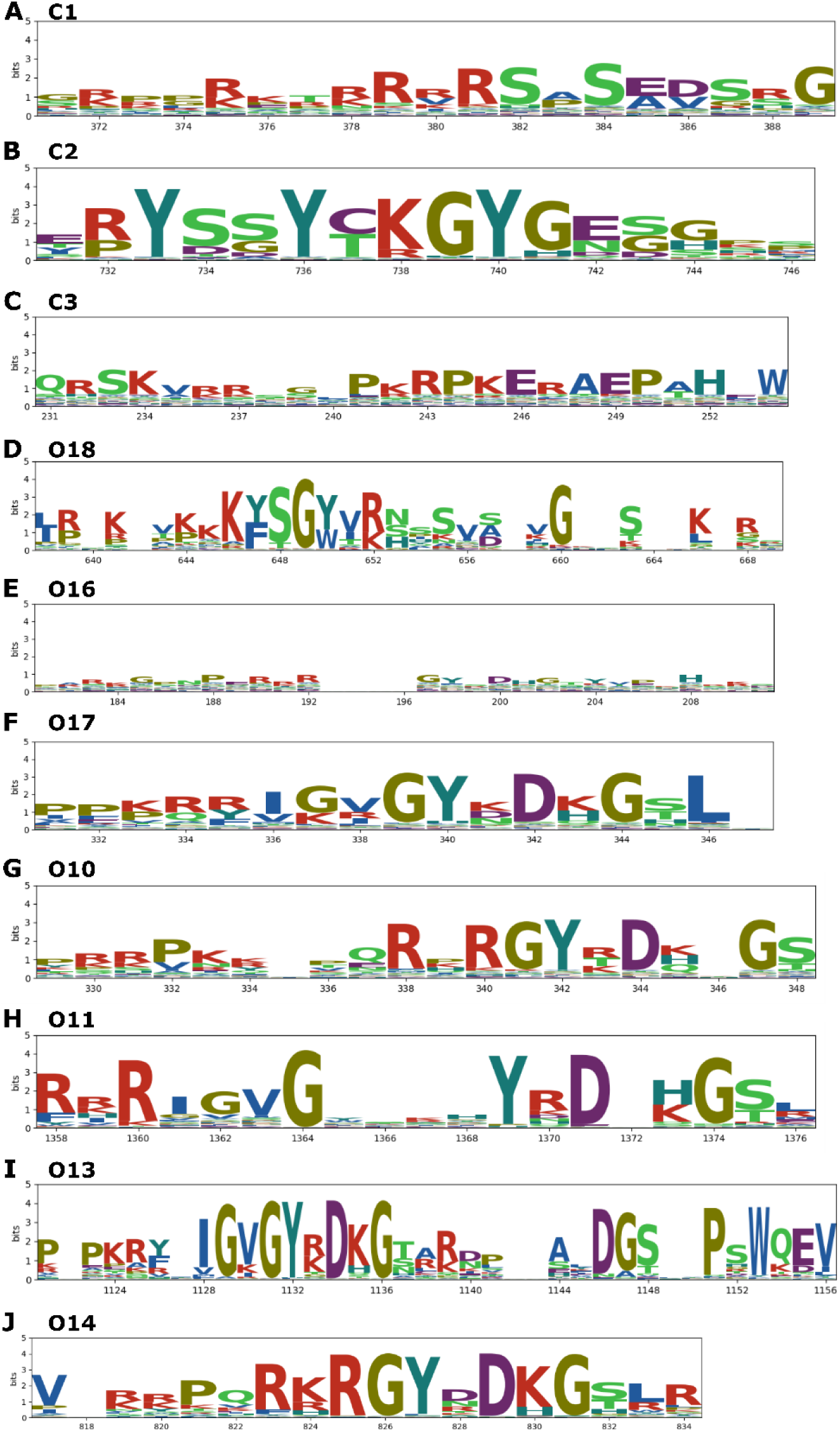
Sequence comparison of the ‘A-domain’ across Oblin-1 subfamilies. (A-J) Stacked sequence logos (information content (bits)) are shown based on the Oblin-1 alignments of the respective subfamilies. A-D show subfamilies lacking the hallmark amino acid signature GYxDxG which is present in the subfamilies shown in E-J.

**Fig. S5:**
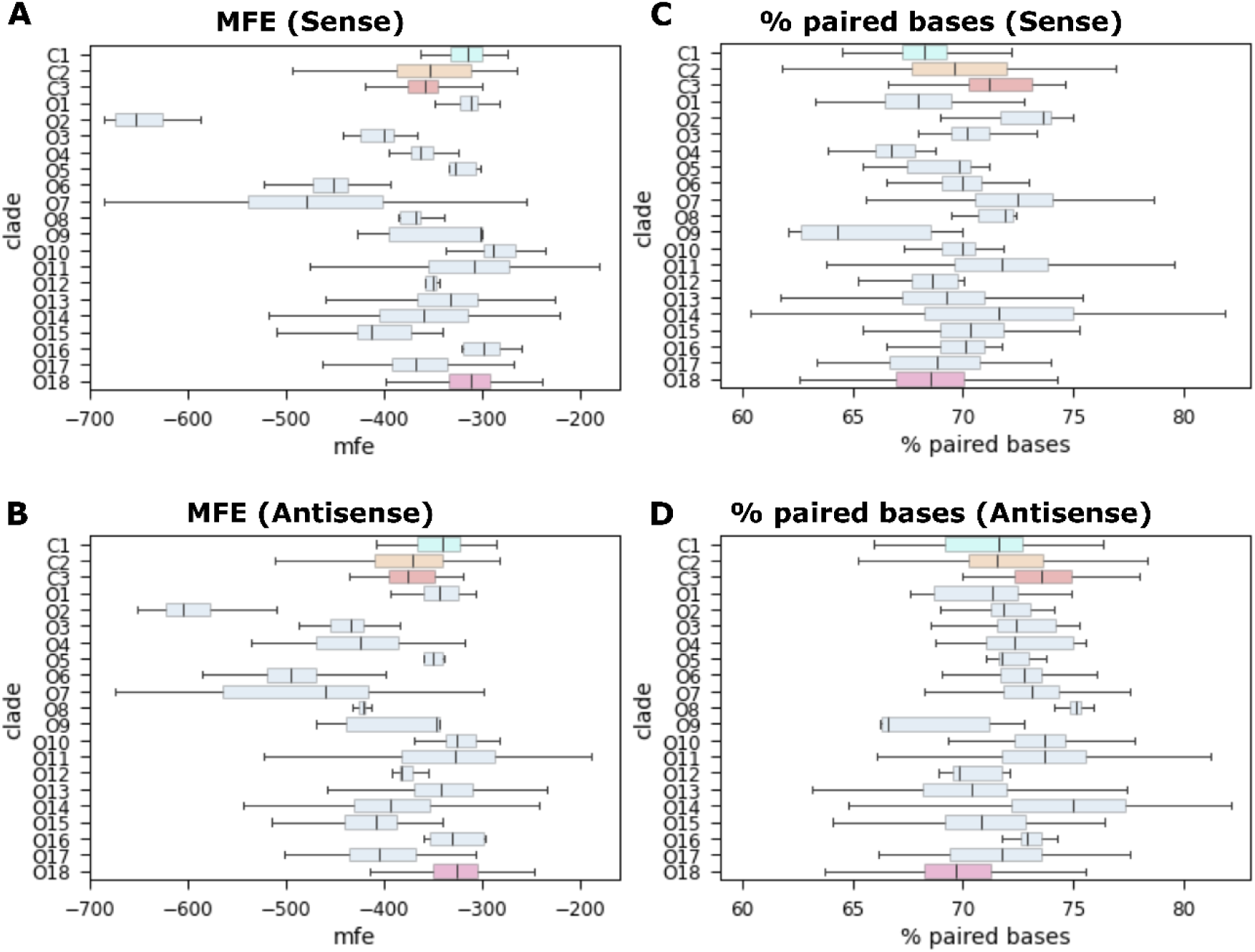
Minimum free energy and percent paired bases of RNA secondary structure prediction. (A,B) Minimum free energy (mfe) and (C,D) percent paired bases for Obelisk RNA secondary structure predictions by RNAfold for both the sense and the antisense genome (with respect to the Oblin-1 ORF).

**Fig. S6:**
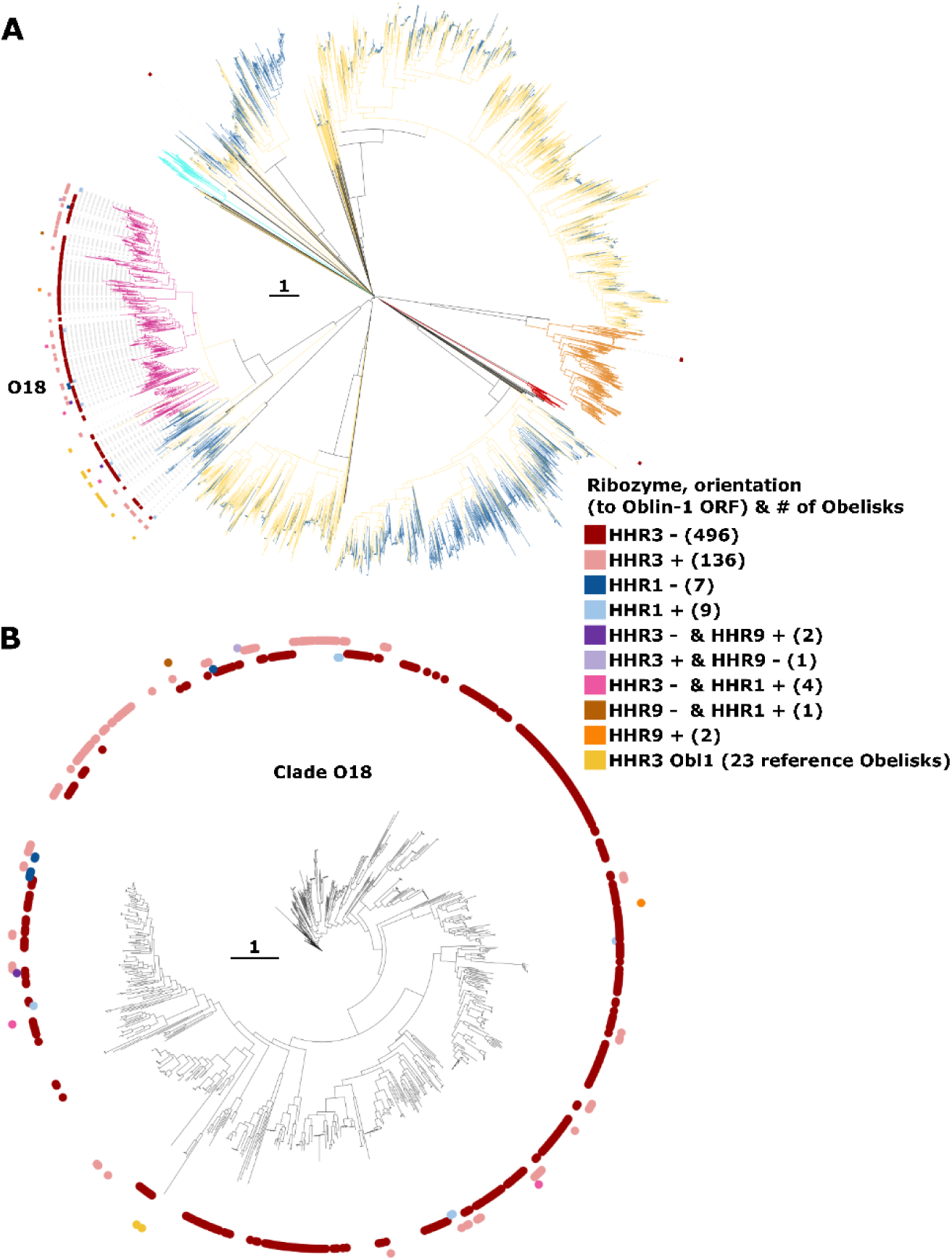
Hammerhead ribozymes across Obelisks. (A) Full Oblin-1 dendrogram as in Fig.3 with Obelisks harboring a hammerhead ribozyme indicated on the rim. (B) Unrooted tree of clade O18 were the rim indicates the presence of hammerhead ribozymes. Legend (for A,B): HHR1, 3, 9: hammerhead ribozyme 1,3 and 9, respectively; ‘+‘ or ‘-’: ribozyme was detected on sense (+) or antisense (-) genome, with respect to the Oblin-1 ORF; HHR3 Obl1: centroid Obelisks from Zheludev et al. (Zheludev et al., 2024) which are reported to have a distinct HHR3; number in brackets gives the number of leaves with this type of ribozyme whereas each leaf can represent multiple Obelisks.

**Fig. S7:**
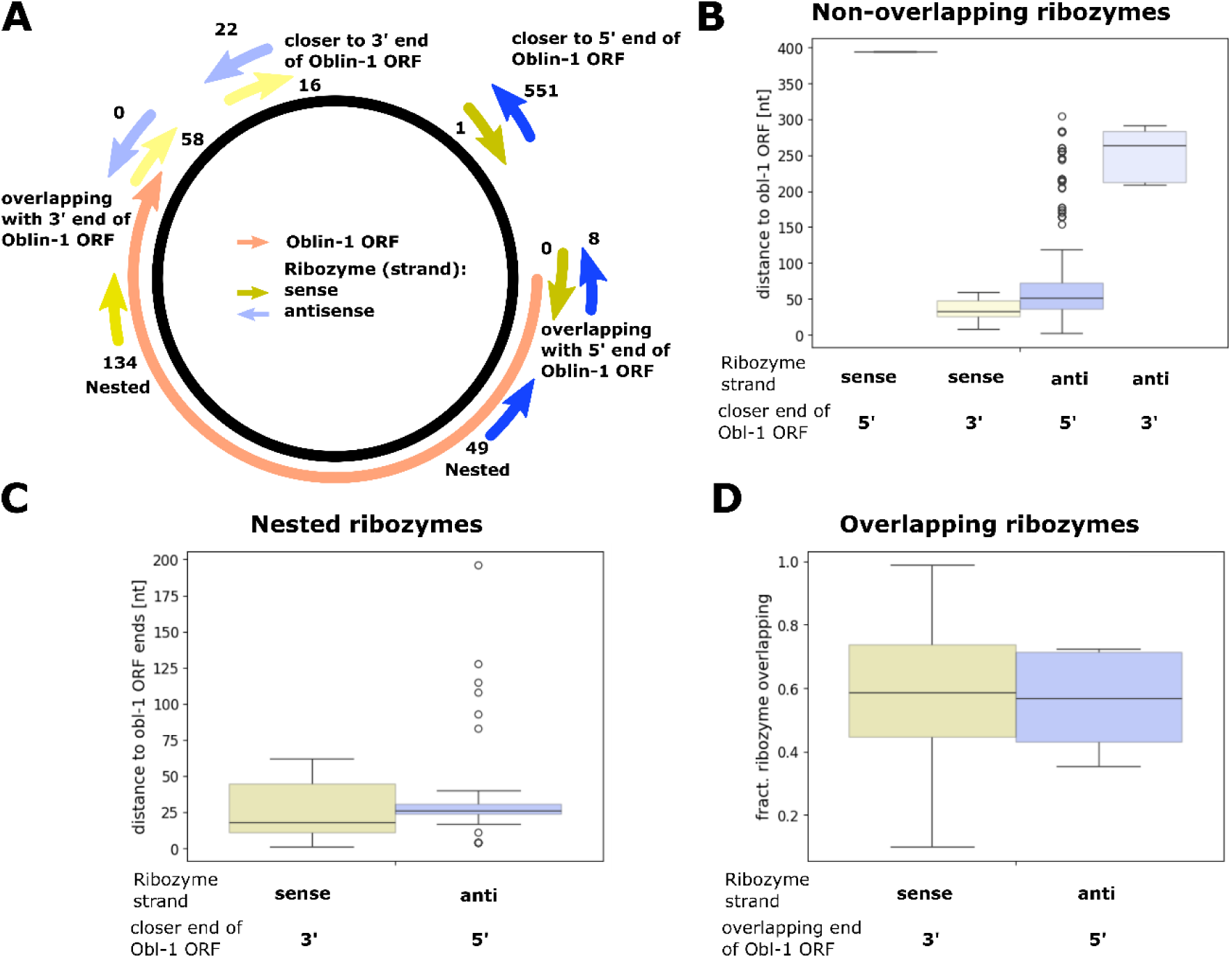
Position of predicted ribozymes. (A) Schematic of the position of the predicted ribozymes and their position in relation to the Oblin-1 ORF on the cccRNA. Numbers indicate observed events of a ribozyme in that position. Arrows in blue indicate ribozymes on the antisense strand compared to the strand encoding the Oblin-1 ORF and arrows in yellow vice versa. (B) Distance of ribozyme to 5’ or 3’ end of Oblin-1 ORF (whichever is closer) if non-overlapping. (C) Distance of nested ribozyme to the 5’ or 3’ end of the Oblin-1 ORF (whichever is closer). No nested ribozymes found in the combination ‘sense and closer to 5’ end’ and ‘antisense and closer to 3’ end’. (D) Fraction of predicted ribozyme overlapping with either the 5’ or the 3’ end of the Oblin-1 ORF.

**Fig. S8:**
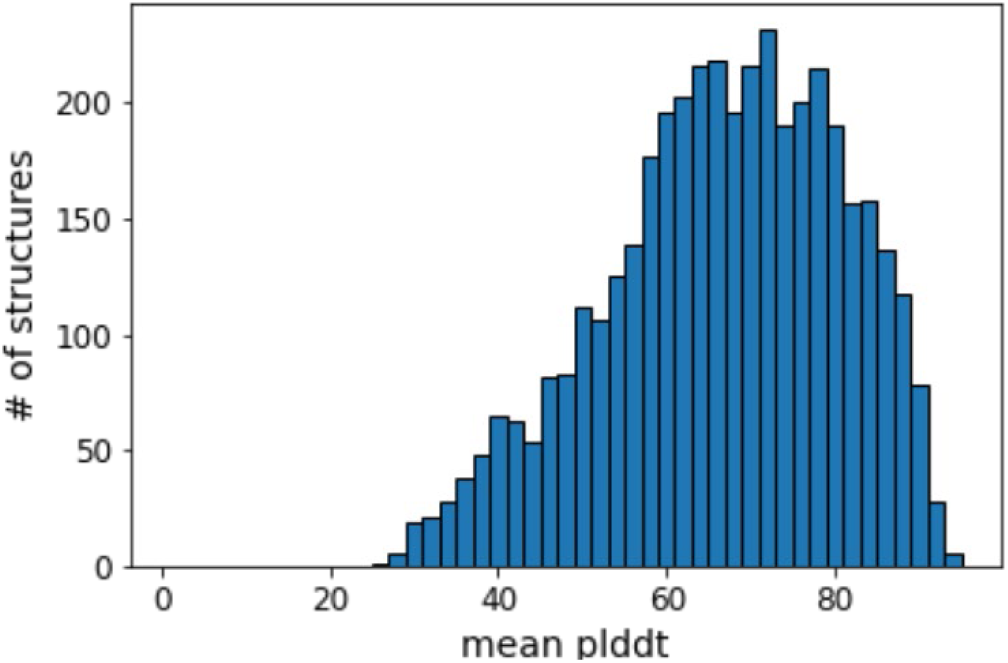
Mean plddt across AlphaFold3 structure predictions for non-Oblin-1 proteins. Histogram of average plddt values for AlphaFold3 predicted structures for non-Oblin-1 proteins.

**Fig. S9:**
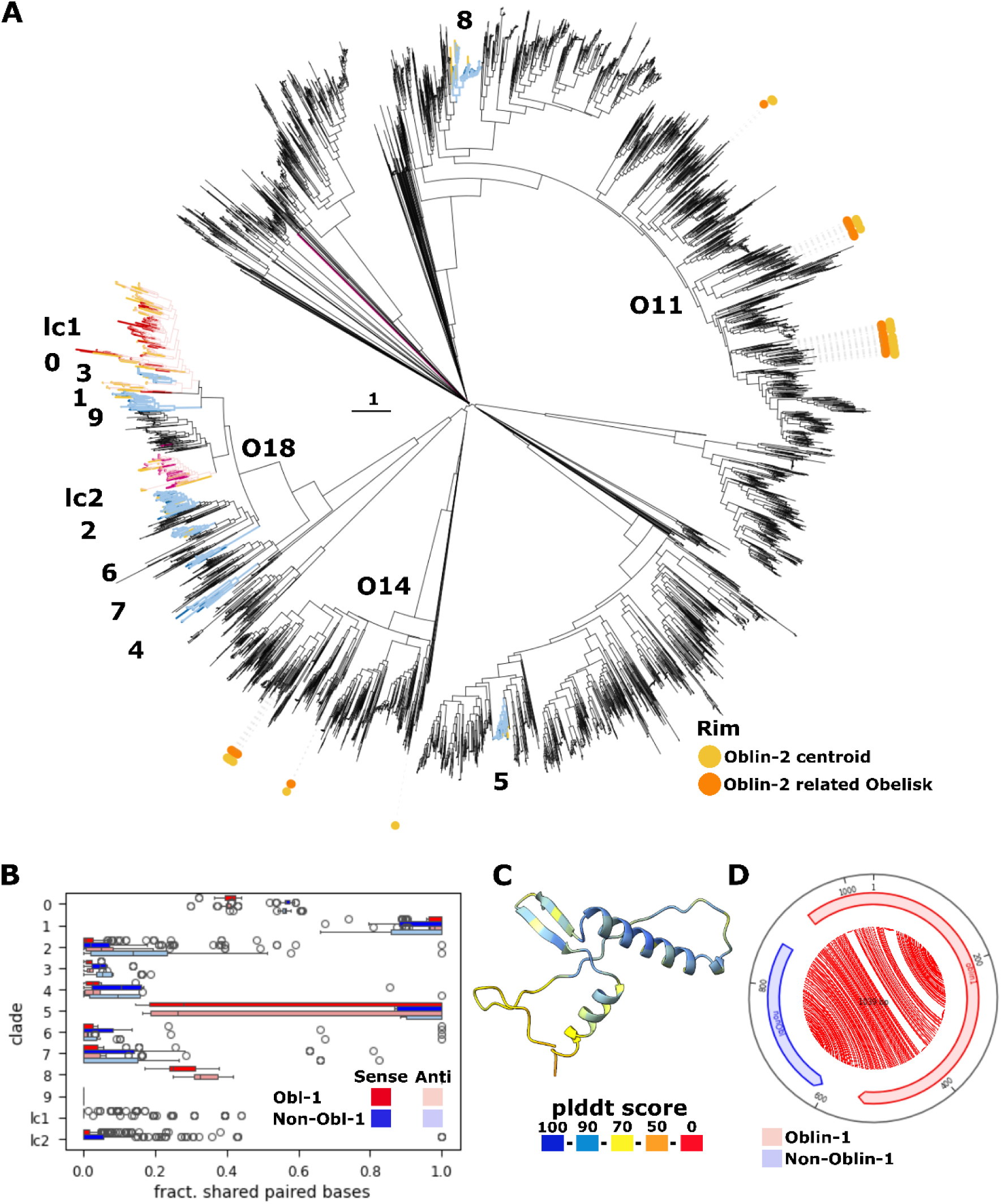
Putative non-Oblin-1 proteins. (A) Clades across the Oblin-1 tree which encode for putative non-Oblin-1 proteins. More than 70% of Obelisks in blue clades harbor the respective non-Oblin-1 protein whereas coverage is lower in red clades (low coverage, lc). Yellow terminals indicate absence of the respective non-Oblin-1 protein in these clades. The presence of Oblin-2 proteins is indicated as yellow and orange dots on the outer rim (reference Obelisks and newly identified ones, respectively). (B) Fraction of paired bases between the non-Oblin-1 (blue) and the Oblin-1 (red) ORF in the RNA secondary structure prediction of the respective Obelisks (both sense (darker color) and antisense (light color) prediction). (C) Representative non-Oblin-1 AlphaFold3 structure prediction from group 2, colored by plddt score. (D) Obelisk map showing the non-Oblin-1 (blue) and the Oblin-1 (red) ORF and a jupyter plot of the paired bases of the structure prediction (RNAfold) of the sense genome. The Obelisks harbors the non-Oblin-1 protein from group 2 shown in C.

**Fig. S10:**
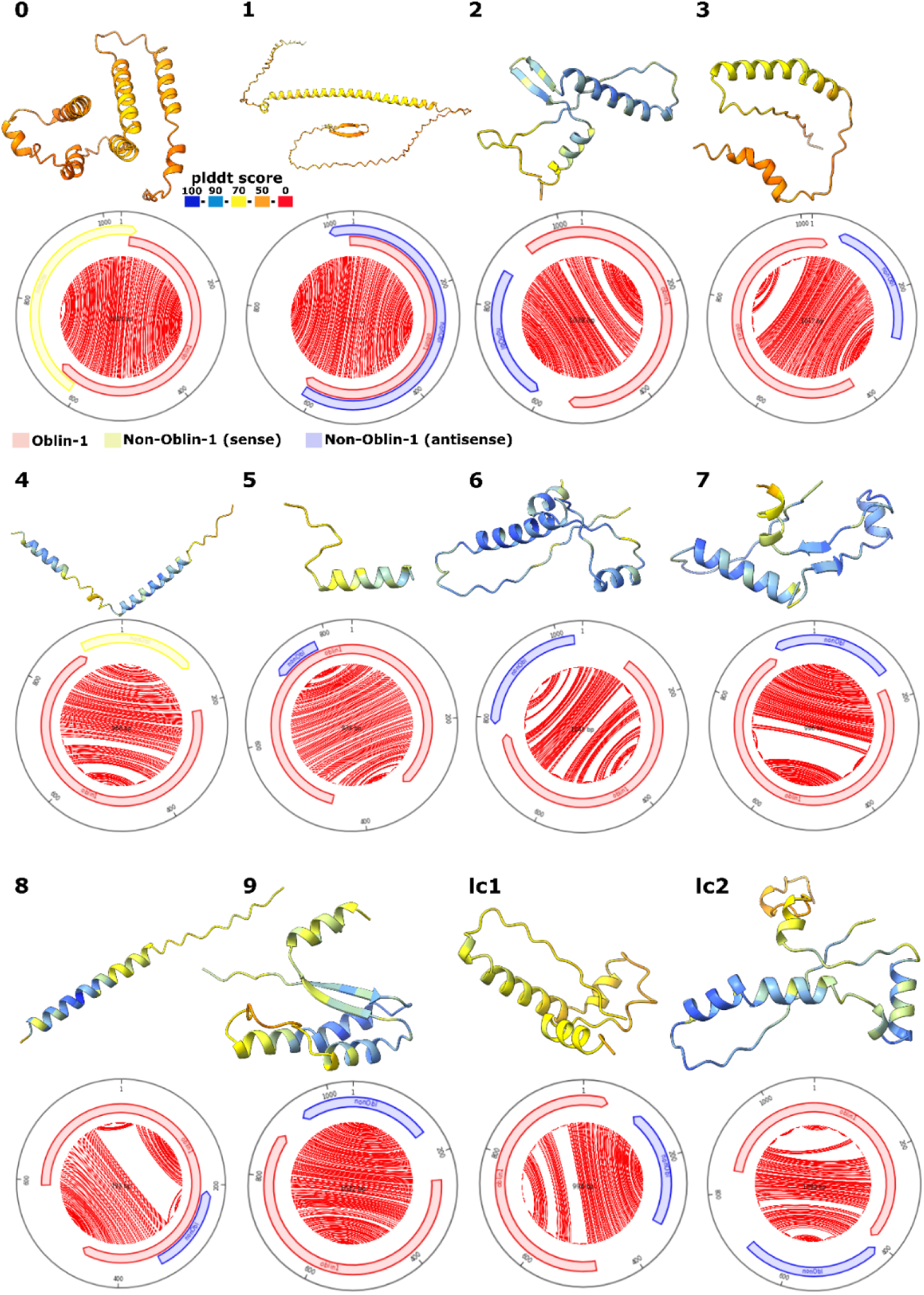
Organization of Obelisks with putative non-Oblin-1 proteins. AlphaFold3 predicted putative non-Oblin-1 structures colored by plddt score from the non-Oblin-1 groups indicated in Fig. S9 (0-9, lc1, lc2). Below each structure is the respective Obelisk map including the respective non-Oblin-1 protein (yellow, if same orientation as Oblin-1 ORF, otherwise blue) and Oblin-1 ORF (red). The jupyter plot of the paired bases of the sense genome of the respective RNA secondary structure prediction (RNAfold) is shown for each map.

## References

Abramson, J., Adler, J., Dunger, J., Evans, R., Green, T., Pritzel, A., Ronneberger, O., Willmore, L., Ballard, A.J., and Bambrick, J. (2024). Accurate structure prediction of biomolecular interactions with AlphaFold 3. Nature 630, 493–500.

Altschul, S.F., Madden, T.L., Schäffer, A.A., Zhang, J., Zhang, Z., Miller, W., and Lipman, D.J. (1997). Gapped BLAST and PSI-BLAST: a new generation of protein database search programs. Nucleic Acids Res 25, 3389–3402.

Bankevich, A., Nurk, S., Antipov, D., Gurevich, A.A., Dvorkin, M., Kulikov, A.S., Lesin, V.M., Nikolenko, S.I., Pham, S., and Prjibelski, A.D. (2012). SPAdes: a new genome assembly algorithm and its applications to single-cell sequencing. J Comput Biol 19, 455–477.

Berman, H.M., Westbrook, J., Feng, Z., Gilliland, G., Bhat, T.N., Weissig, H., Shindyalov, I.N., and Bourne, P.E. (2000). The protein data bank. Nucleic Acids Res 28, 235–242.

Branch, A.D., and Robertson, H.D. (1984). A replication cycle for viroids and other small infectious RNA’s. Science 223, 450–455.

Chandonia, J.-M., Guan, L., Lin, S., Yu, C., Fox, N.K., and Brenner, S.E. (2022). SCOPe: improvements to the structural classification of proteins–extended database to facilitate variant interpretation and machine learning. Nucleic Acids Res 50, D553–D559.

Chen, I.-M.A., Chu, K., Palaniappan, K., Ratner, A., Huang, J., Huntemann, M., Hajek, P., Ritter, S., Varghese, N., and Seshadri, R. (2021). The IMG/M data management and analysis system v. 6.0: new tools and advanced capabilities. Nucleic Acids Res 49, D751–D763.

Cheng, H., Schaeffer, R.D., Liao, Y., Kinch, L.N., Pei, J., Shi, S., Kim, B.-H., and Grishin, N.V. (2014). ECOD: an evolutionary classification of protein domains. PLoS Comp Biol 10, e1003926.

Chong, L.C., and Lauber, C. (2023). Viroid-like RNA-dependent RNA polymerase-encoding ambiviruses are abundant in complex fungi. Front Microbiol 14, 1144003.

Di Serio, F., Flores, R., Verhoeven, J.T.J., Li, S.-F., Pallás, V., Randles, J., Sano, T., Vidalakis, G., and Owens, R. (2014). Current status of viroid taxonomy. Arch Virol 159, 3467–3478.

Di Serio, F., Li, S.-F., Matoušek, J., Owens, R.A., Pallás, V., Randles, J.W., Sano, T., Verhoeven, J.T.J., Vidalakis, G., and Flores, R. (2018). ICTV virus taxonomy profile: Avsunviroidae. J Gen Virol 99, 611–612.

Di Serio, F., Owens, R.A., Li, S.-F., Matoušek, J., Pallás, V., Randles, J.W., Sano, T., Verhoeven, J.T.J., Vidalakis, G., and Flores, R. (2021). ICTV virus taxonomy profile: Pospiviroidae. J Gen Virol 102, 001543.

Edgar, R.C. (2022). Muscle5: High-accuracy alignment ensembles enable unbiased assessments of sequence homology and phylogeny. Nature communications 13, 6968.

Fall, M.L., Xu, D., Lemoyne, P., Moussa, I.E.B., Beaulieu, C., and Carisse, O. (2020). A Diverse Virome of Leafroll-Infected Grapevine Unveiled by dsRNA Sequencing. Viruses 12, 1142.

Flores, R., Gas, M.E., Molina-Serrano, D., Nohales, M.A., Carbonell, A., Gago, S., De la Pena, M., and Daros, J.A. (2009). Viroid replication: rolling-circles, enzymes and ribozymes. Viruses 1, 317–334.

Forgia, M., Navarro, B., Daghino, S., Cervera, A., Gisel, A., Perotto, S., Aghayeva, D.N., Akinyuwa, M.F., Gobbi, E., Zheludev, I.N., et al. (2023). Hybrids of RNA viruses and viroid-like elements replicate in fungi. Nat Commun 14, 2591.

Fox, N.K., Brenner, S.E., and Chandonia, J.-M. (2014). SCOPe: Structural Classification of Proteins—extended, integrating SCOP and ASTRAL data and classification of new structures. Nucleic Acids Res 42, D304–D309.

Gruber, A.R., Lorenz, R., Bernhart, S.H., Neuböck, R., and Hofacker, I.L. (2008). The vienna RNA websuite. Nucleic Acids Res 36, W70–W74.

Hirai, M., Takaki, Y., Kondo, F., Horie, M., Urayama, S., and Nunoura, T. (2021). RNA Viral Metagenome Analysis of Subnanogram dsRNA Using Fragmented and Primer Ligated dsRNA Sequencing (FLDS). Microbes Environ 36, ME20152.

Holm, L. (2022). Dali server: structural unification of protein families. Nucleic Acids Res 50, W210–W215.

Jumper, J., Evans, R., Pritzel, A., Green, T., Figurnov, M., Ronneberger, O., Tunyasuvunakool, K., Bates, R., Žídek, A., and Potapenko, A. (2021). Highly accurate protein structure prediction with AlphaFold. Nature 596, 583–589.

Koonin, E.V. (2024). Circular RNAs from linear viral RNA genomes: A distinct dimension in the virus world. Proc Natl Acad Sci U S A 121, e2401335121.

Koonin, E.V., Kuhn, J.H., Dolja, V.V., and Krupovic, M. (2024). Megataxonomy and global ecology of the virosphere. ISME J 18, wrad042.

Koonin, E.V., and Lee, B.D. (2025). Diversity and evolution of viroids and viroid-like agents with circular RNA genomes revealed by metatranscriptome mining. Nucleic Acids Res 53, gkae1278.

Kuhn, J.H., Babaian, A., Bergner, L.M., Dény, P., Glebe, D., Horie, M., Koonin, E.V., Krupovic, M., Paraskevopoulou, S., and de la Peña, M. (2024a). ICTV virus taxonomy profile: Kolmioviridae 2024. J Gen Virol 105, 001963.

Kuhn, J.H., Botella, L., de la Peña, M., Vainio, E.J., Krupovic, M., Lee, B.D., Navarro, B., Sabanadzovic, S., Simmonds, P., and Turina, M. (2024b). Ambiviricota, a novel ribovirian phylum for viruses with viroid-like properties. J Virol 98, e00831–00824.

Lasda, E., and Parker, R. (2014). Circular RNAs: diversity of form and function. RNA 20, 1829–1842.

Lee, B.D., and Koonin, E.V. (2022). Viroids and Viroid-like Circular RNAs: Do They Descend from Primordial Replicators? Life (Basel) 12, 103.

Lee, B.D., Neri, U., Oh, C.J., Simmonds, P., and Koonin, E.V. (2022). ViroidDB: a database of viroids and viroid-like circular RNAs. Nucleic Acids Res 50, D432–D438.

Lee, B.D., Neri, U., Roux, S., Wolf, Y.I., Camargo, A.P., Krupovic, M., Consortium, R.N.A.V.D., Simmonds, P., Kyrpides, N., Gophna, U., et al. (2023). Mining metatranscriptomes reveals a vast world of viroid-like circular RNAs. Cell 186, 646–661 e644.

Letunic, I., and Bork, P. (2024). Interactive Tree of Life (iTOL) v6: recent updates to the phylogenetic tree display and annotation tool. Nucleic Acids Res 52, W78–W82.

Liu, C.X., and Chen, L.L. (2022). Circular RNAs: Characterization, cellular roles, and applications. Cell 185, 2016–2034.

Lopez-Simon, J., de la Pena, M., and Martinez-Garcia, M. (2025). Viroid-like “obelisk” agents are widespread in the ocean and exceed the abundance of RNA viruses in the prokaryotic fraction. ISME J 19, wraf033.

Lorenz, R., Bernhart, S.H., Höner zu Siederdissen, C., Tafer, H., Flamm, C., Stadler, P.F., and Hofacker, I.L. (2011). ViennaRNA Package 2.0. Algorithms Mol Biol 6, 1–14.

Medvedeva, S., Liu, Y., Koonin, E.V., Severinov, K., Prangishvili, D., and Krupovic, M. (2019). Virus-borne mini-CRISPR arrays are involved in interviral conflicts. Nature communications 10, 5204.

Mirdita, M., Schütze, K., Moriwaki, Y., Heo, L., Ovchinnikov, S., and Steinegger, M. (2022). ColabFold: making protein folding accessible to all. Nat Methods 19, 679–682.

Mistry, J., Chuguransky, S., Williams, L., Qureshi, M., Salazar, G.A., Sonnhammer, E.L., Tosatto, S.C., Paladin, L., Raj, S., and Richardson, L.J. (2021). Pfam: The protein families database in 2021. Nucleic Acids Res 49, D412–D419.

Morris, T.J., and Dodds, J.A. (1979). Isolation and analysis of double-stranded-RNA from virus-infected plant and fungal tissue. Phytopathology 69, 854–858.

Navarro, B., and Turina, M. (2024). Viroid and viroid-like elements in plants and plant-associated microbiota: a new layer of biodiversity for plant holobionts. New Phytol 244, 1216–1222.

Nawrocki, E.P., and Eddy, S.R. (2013). Infernal 1.1: 100-fold faster RNA homology searches. Bioinformatics 29, 2933–2935.

Neri, U., Wolf, Y.I., Roux, S., Camargo, A.P., Lee, B., Kazlauskas, D., Chen, I.M., Ivanova, N., Allen, L.Z., and Paez-Espino, D. (2022). Expansion of the global RNA virome reveals diverse clades of bacteriophages. Cell 185, 4023–4037.

Nishimura, Y., and Yoshizawa, S. (2022). The OceanDNA MAG catalog contains over 50,000 prokaryotic genomes originated from various marine environments. Scientific Data 9, 305.

Pettersen, E.F., Goddard, T.D., Huang, C.C., Meng, E.C., Couch, G.S., Croll, T.I., Morris, J.H., and Ferrin, T.E. (2021). UCSF ChimeraX: Structure visualization for researchers, educators, and developers. Protein Sci 30, 70–82.

Price, M.N., Dehal, P.S., and Arkin, A.P. (2009). FastTree: computing large minimum evolution trees with profiles instead of a distance matrix. Mol Biol Evol 26, 1641–1650.

Price, M.N., Dehal, P.S., and Arkin, A.P. (2010). FastTree 2–approximately maximum-likelihood trees for large alignments. PLoS One 5, e9490.

Roossinck, M.J., Saha, P., Wiley, G.B., Quan, J., White, J.D., Lai, H., Chavarria, F., Shen, G., and Roe, B.A. (2010). Ecogenomics: using massively parallel pyrosequencing to understand virus ecology. Mol Ecol 19 Suppl 1, 81–88.

Söding, J. (2005). Protein homology detection by HMM–HMM comparison. Bioinformatics, 951–960.

Semancik, J.S. (1986). Separation of viroid RNAs by cellulose chromatography indicating conformational distinctions. Virology 155, 39–45.

Steinegger, M., Meier, M., Mirdita, M., Vohringer, H., Haunsberger, S.J., and Soding, J. (2019). HH-suite3 for fast remote homology detection and deep protein annotation. BMC Bioinformatics 20, 473.

Steinegger, M., and Söding, J. (2017). MMseqs2 enables sensitive protein sequence searching for the analysis of massive data sets. Nat Biotechnol 35, 1026–1028.

Tareen, A., and Kinney, J.B. (2020). Logomaker: beautiful sequence logos in Python. Bioinformatics 36, 2272–2274.

Urayama, S., Fukudome, A., Hirai, M., Okumura, T., Nishimura, Y., Takaki, Y., Kurosawa, N., Koonin, E.V., Krupovic, M., and Nunoura, T. (2024). Double-stranded RNA sequencing reveals distinct riboviruses associated with thermoacidophilic bacteria from hot springs in Japan. Nat Microbiol 9, 514–523.

Urayama, S., Takaki, Y., and Nunoura, T. (2016). FLDS: A comprehensive dsRNA sequencing method for intracellular RNA virus surveillance. Microbes Environ 31, 33–40.

Van Kempen, M., Kim, S.S., Tumescheit, C., Mirdita, M., Lee, J., Gilchrist, C.L., Söding, J., and Steinegger, M. (2024). Fast and accurate protein structure search with Foldseek. Nat Biotechnol 42, 243–246.

Wolf, Y.I., Kazlauskas, D., Iranzo, J., Lucia-Sanz, A., Kuhn, J.H., Krupovic, M., Dolja, V.V., and Koonin, E.V. (2018). Origins and Evolution of the Global RNA Virome. mBio 9, e02329–02318.

Wolf, Y.I., Silas, S., Wang, Y., Wu, S., Bocek, M., Kazlauskas, D., Krupovic, M., Fire, A., Dolja, V.V., and Koonin, E.V. (2020). Doubling of the known set of RNA viruses by metagenomic analysis of an aquatic virome. Nature microbiology 5, 1262–1270.

Zheludev, I.N., Edgar, R.C., Lopez-Galiano, M.J., de la Pena, M., Babaian, A., Bhatt, A.S., and Fire, A.Z. (2024). Viroid-like colonists of human microbiomes. Cell 187, 6521–6536 e6518.

